# Rare “Jackpot” Individuals Drive Rapid Adaptation in Threespine Stickleback

**DOI:** 10.1101/2025.03.25.642177

**Authors:** Alexander Kwakye, Kerry Reid, Matthew A. Wund, David C. Heins, Michael A. Bell, Krishna R. Veeramah

## Abstract

Recombination has long been considered the primary mechanism to bring beneficial alleles together, which can increase the speed of adaptation from standing genetic variation. Recombination is fundamental to the transporter hypothesis proposed to explain precise parallel adaptation in Threespine Stickleback. We studied an instance of freshwater adaptation in the Threespine Stickleback system using whole genomic data from an evolutionary time series to observe the genomic dynamics underlying rapid parallel adaptation. Our experiment showed that rapid adaptation to a freshwater environment depended on a few individuals with large haploblocks of freshwater-adaptive alleles (jackpot carriers) present among the anadromous (i.e., sea-run) founders at low frequencies. Biological kinship analyses indicate that mating among jackpot carriers and between jackpot carriers and non-jackpot individuals led to a rapid increase in freshwater-adaptive alleles within the first few generations. This process allowed the population to overcome a substantial bottleneck likely caused by the low fitness of first-generation stickleback with a few freshwater-adaptive alleles born in the lake. Additionally, we found evidence that the genetic load that emerged from population growth after the bottleneck may have been reduced through an increase in homozygosity by inbreeding, ultimately purging deleterious alleles. Recombination likely played a limited role in this case of very rapid adaptation.

## INTRODUCTION

Darwin originally proposed that evolution by natural selection is a gradual process whereby changes in phenotypes that led to speciation occur by selection of variants with small fitness differences over millions of years^1^. However, there are now numerous examples of rapid evolution, i.e., within tens of generations^2–5^, which provide an opportunity to study the molecular basis of evolution in more complex eukaryotes outside of a lab setting or during selective breeding of domesticated species.

Rapid adaptation proceeds predominantly from standing genetic variation (SGV)^6–10^, and *Gasterosreus aculeatus* (i.e., Threespine Stickleback fish) is a premier example of this phenomenon^11^. In this nominal species, hundreds of loci, most estimated to be more than a million years old^12^, are the basis for much more recent, precisely repeated, rapid adaptation of ancestral marine or anadromous (i.e., collectively ‘oceanic’) ecotypes to freshwater habitats^12–14^. This process has been shown to occur within a decade^12,15^.

According to the transporter hypothesis^16^ (see also ref ^10^), this ancient SGV is maintained at low frequencies in oceanic populations through persistent gene flow from multiple freshwater populations, which are sympatric with anadromous populations during the breeding season. Oceanic-freshwater hybrids with haploblocks (large chromosomal segments with multiple contiguous loci) of freshwater-adaptive alleles move back into the oceanic populations, where they should be disfavored but survive well enough to backcross to anadromous stickleback. Successive generations of backcrossing with oceanic stickleback and recombination break these blocks of freshwater-adaptive alleles apart, and most oceanic individuals are observed to possess only a small number of them, mostly in the heterozygous state^12^. Consequently, adaptation to new freshwater habitats is thought to proceed from colonizing oceanic individuals, each possessing only a few freshwater-adaptive alleles. In freshwater habitats, selection is then believed to favor the freshwater-adaptive alleles, leading to an increase in their frequencies, and genetic linkage among these alleles will allow for their re-assembly back into large haploblocks over multiple generations.

Alternatively, rapid freshwater adaptation may involve selection that strongly favors jackpot carriers. Jackpot carriers are rare oceanic individuals that possess large haploblocks of freshwater-adaptive alleles. Previous studies suggest that the frequency of jackpot carriers in marine environments is about 0.1%^12,17^. Bassham et al.^17^ proposed that rapid freshwater adaptation may depend on such jackpot carriers. In addition, simulations have found that a few individuals with large haploblocks of freshwater-adaptive alleles within founding oceanic populations would be sufficient to adapt to new freshwater habitats^18^. The presence of such individuals could dramatically increase the speed with which haplotypes of adaptive alleles spread through the population compared to the waiting time required for recombination to bring these adaptive alleles together within individuals. In addition, previous work has shown that recombination is significantly suppressed in regions of the genome containing freshwater-adaptive alleles, allowing the maintenance of what has been termed “adaptive cassettes”^12,19,20^.

Evolutionary experiments that introduce oceanic Threespine Stickleback into new freshwater habitats^21–23^ allow us to examine the freshwater-adaptive process forward-in-time rather than inferring it retrospectively. This approach provides a way to test these two hypotheses of how Threespine Stickleback adapt to new freshwater habitats: reassembly through recombination versus a jackpot carrier-mediated adaptive process.

In this study, we examined a recently founded freshwater population in Scout Lake, Alaska, derived from anadromous ancestors^21^. We sequenced hundreds of whole genomes from samples collected during the first few generations of rapid adaptation to conditions in Scout Lake. Our results show that adaptation to the freshwater environment depended on the presence of a few jackpot carriers among the founders. After a bottleneck detected by both genomic analyses and catch per unit effort (CPUE), population growth was driven primarily by breeding among the descendants of those few jackpot carriers, leaving signals of marked inbreeding in subsequent generations. The increase in homozygosity resulting from inbreeding facilitated the removal of deleterious alleles, thereby likely improving the population’s adaptability. Our empirical study shows that only a few jackpot carriers can drive rapid adaptation of anadromous Threespine Stickleback to new freshwater habitats. These findings also suggest a limited role of recombination during the earliest stages of rapid adaptation, in contrast with predictions from the transporter hypothesis.

## RESULTS

### The Scout Lake experiment

Scout Lake (60.5353N, 150.8322W) is on the Kenai Peninsula, Alaska, USA. After treatment with rotenone in 2009 to exterminate an invasive fish species, 3047 anadromous Threespine Stickleback from a spawning run in Rabbit Slough (62.6893N, 149.427E) in the Matanuska-Susitna Borough, Alaska, were released into the lake in the summer of 2011 (fig. 1A). Tests with trout late in the summer of 2009 indicated that fish did not survive the rotenone treatment (Supplementary Section 1). The lake is isolated from other bodies of water, and sticklebacks (and other fish) were not detected by exhaustive sampling. Stickleback samples have been collected annually from this lake since the year after its founding in 2011 (i.e. from 2012) for phenotypic and genetic analysis^12,21^. In this study, we focused on samples collected in 2013, 2014, 2015, 2017 and 2020 and we referred to these samples as SC2013, SC2014, SC2015, SC2017 and SC2020, respectively. These samples correspond to 2, 3, 4, 6 and 9 years since the founding of Scout Lake. In addition, a sample from the anadromous source population, Rabbit Slough, is denoted as RS2019.

**Fig. 1:**
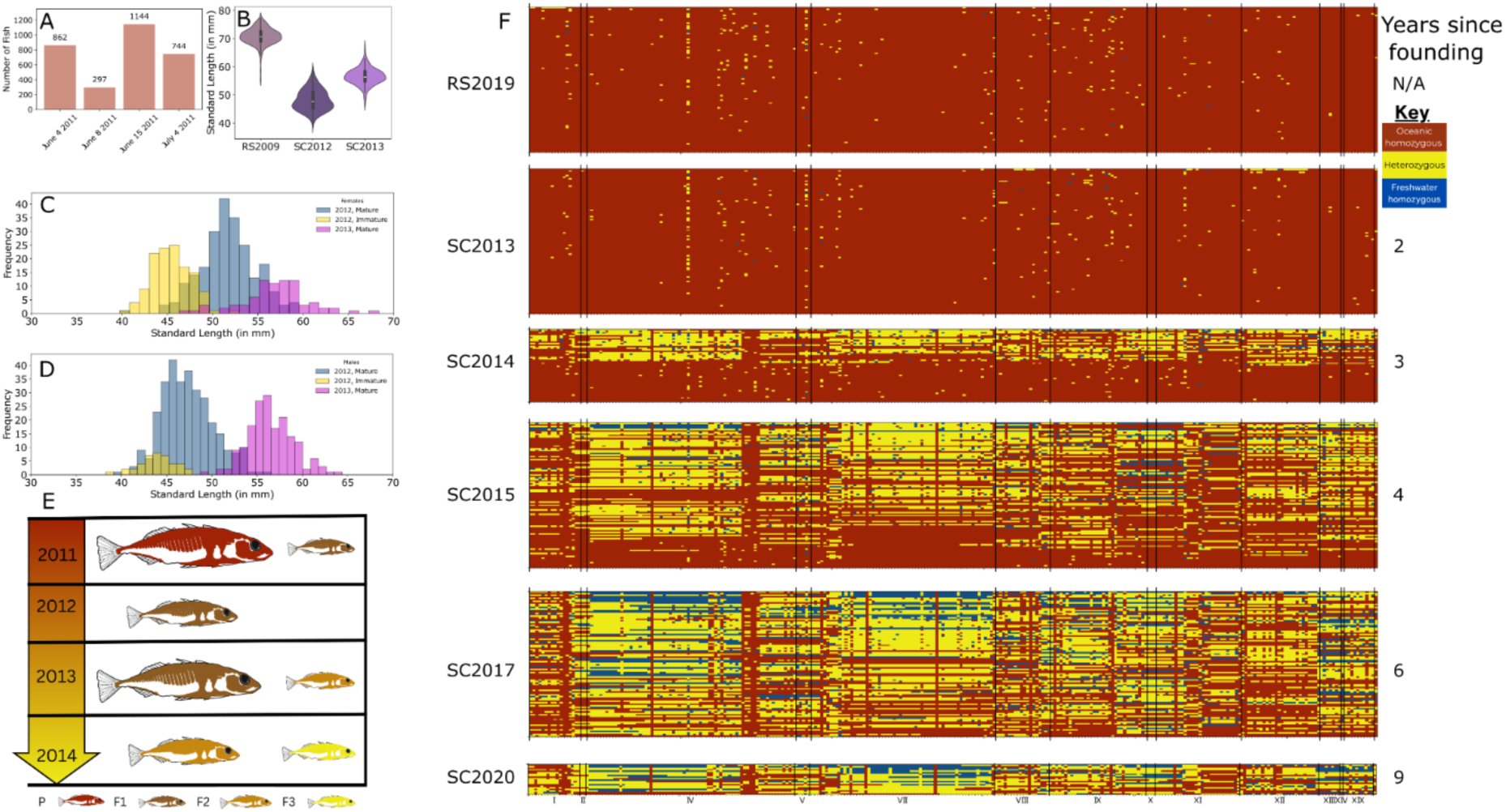
Genotypes of freshwater adaptive loci: (**A).** Number of anadromous Rabbit Slough used to found Scout Lake in 2011. (**B).** Standard length (SL) of Rabbit Slough 2009, SC2012 and SC2013 samples.The SL is a measure of the size of the fish. **(C).** Sexual maturity of females in SC2012 and SC2013 samples. (**D).** Same as C but for males. (**E).** Inferred generations based on panels B, C and D for the samples collected in the first three years after founding of Scout Lake. (**F).** Genotypes of individuals used in this study at freshwater adaptive loci. Each column is a multi-SNP locus and each row is an individual genome. N/A years since founding indicates specimens collected in Rabbit Slough in 2019, a representative of the anadromous founders of Scout Lake stickleback population; 2,3,4,6 and 9 years since founding are samples in Scout Lake collected in 2013, 2014, 2015, 2017 and 2020 respectively. We denote samples from 2013, 2014, 2015, 2017 and 2020 as SC2013,SC2014, SC2015,SC2017, and SC2020 respectively. The Rabbit Slough sample is denoted by RS2019. There are 96 specimens in RS2019, SC2013, SC2015 and SC2017 samples; 47 and 20 in SC2014 and SC2020 samples respectively. Each locus has three to 3658 SNPs. For B and C, the bars have three colors: yellow, light blue and purple. The other colors result from overlap of these colors.

The adults stocked in Scout Lake in 2011 produced offspring after release but did not survive the winter (fig. 1B and Fig. 3 from Kurz et al.^24^). The F1 progeny likely did not successfully reproduce until 2013. However, small sexually mature males and females were captured in 2012 (fig. 1B and 1C, Kurtz et al.^24^ and Supplementary Section 2). This finding indicates that the F1 survived in the lake for two years, which is not surprising given that F1 offspring of anadromous Threespine Stickleback typically spend part of the earliest and latest segments of their life cycle in freshwater^25,26^ and thus are physiologically capable of tolerating freshwater. These phenotypic results show that the 2014 sample (SC2014) represents one-year-old F2 offspring of the F1 generation (fig. 1D), as any stickleback born in the sampling year would have been too small to be captured with the sampling methods employed (i.e. minnow traps). Thus, the first generation of offspring of the parents born and that spent their entire life in the lake were adults in 2014.

### Jackpot carriers increased in frequency in Scout Lake

To infer whether haplotypes with numerous freshwater-adaptive alleles were assembled after introduction to the lake or were already present in rare large haploblocks in a few of the introduced anadromous founders, we analyzed whole genomes from samples collected during the first nine years of founding of Scout Lake and a representative sample from the anadromous Rabbit Slough (RS2019) population used to seed it. We used a TN5 transposase-based approach to sequence 432 genomes at coverages ranging from 0.43X to 1.84X (low-coverage). There were 336 genomes from individuals collected from two (n = 96), three (n = 48), four (n = 96) and six (n = 96) years after the Scout Lake population was founded and 96 genomes from Rabbit Slough. We also sequenced 20 genomes from a sample collected nine years after Scout Lake was founded at a mean of 25.7X coverage (high-coverage) (Methods, Supplementary Section 5, Supplementary Table 1).

We defined multi-SNP haplotypes for each of 344 loci with freshwater-adaptive alleles that we previously identified to experience rapid and significant frequency increases in three lake stickleback populations from Cook Inlet, Alaska^12,22^. These populations, which included the Scout Lake population, were founded by anadromous ancestors. The 344 loci range from a few base pairs to kilobases, with a median size of 27.3 kb. Each multi-SNP haplotype consists of the most significant freshwater-adaptive SNP within the locus and selected neighboring SNPs whose freshwater allele frequencies are highly correlated (r> =0.99) across our time-series data from the three Threespine Stickleback populations^12^. We excluded loci with less than three SNPs, leaving 300 haplotypes that contain three to 3658 SNPs.

We then developed an approximate genotype likelihood approach suitable to call the diploid state of freshwater-adaptive loci as homozygous oceanic, heterozygous, or homozygous freshwater from low-coverage genomic data (Supplementary Section 6). We validated this method by applying it to experimental crosses between parents with known oceanic and freshwater ecotypes derived from a lake (60.914517°, −149.101545°) at mile 87 of Seward Highway a few kilometers east of Girdwood, Alaska (Supplementary Section 4). Genotypic states at loci inferred using our likelihood method were consistent with expectations from our crosses (fig. S1). Parents were scored as homozygous for the alleles associated with their ecotype at most loci. More importantly, the offspring of parents with alternative homozygous states for a locus were heterozygous at that locus (Supplementary Section 6, fig. S1).

Freshwater-adaptive loci were almost always homozygous for oceanic alleles in the ancestral RS2019 and SC2013 (2 years since founding) samples (fig. 1F). Individuals in these samples were either completely homozygous for oceanic alleles (12 and 15 out of 96 specimens for RS2019 and SC2013, respectively) or possessed freshwater-adaptive alleles at only a few loci (average number of freshwater-adaptive alleles in RS2019 = 2.1 [range: 0-9], SC2013 = 2.25 [range: 0-30]). The mean proportion of freshwater-adaptive alleles did not differ significantly between these two samples (SC2013 = 0.0048 vs. RS2019 = 0.0045; *t* = 0.395, df = 134.71, *p-value* = 0.6932, fig.1, S2). Out of the 81 individuals in SC2013 (mostly F1 generation in the lake) with freshwater-adaptive alleles, only 7 of them had two or more loci forming a contiguous block with freshwater-adaptive alleles, with blocks of alleles spanning between 0.17Mb to 3.86Mb [0.41cM to 4.57cM] (fig. 2A, Supplementary table 2). We estimated the size of a contiguous block of freshwater-adaptive alleles by the genome coordinates that defined SNPs. For a series of adjacent loci with freshwater-adaptive alleles, we estimated the size of the block by taking the difference between the first SNP of the first locus with a freshwater-adaptive allele and the last SNP of that series. Only one of the SC2013 specimens possessed notable contiguous blocks of freshwater-adaptive alleles on multiple chromosomes. This individual genome contained a 0.98Mb [0.47cM] block of three heterozygous loci on chrVIII; 3.4Mb [2.95cM] of two heterozygous and one freshwater homozygous loci on chrXIII; nine heterozygous and one homozygous freshwater loci of size 3.86Mb[4.57cM] on chrXX; and eight heterozygous loci totalling 2.43Mb [1.99cM] on chrXX (Supplementary table 2).

**Fig. 2:**
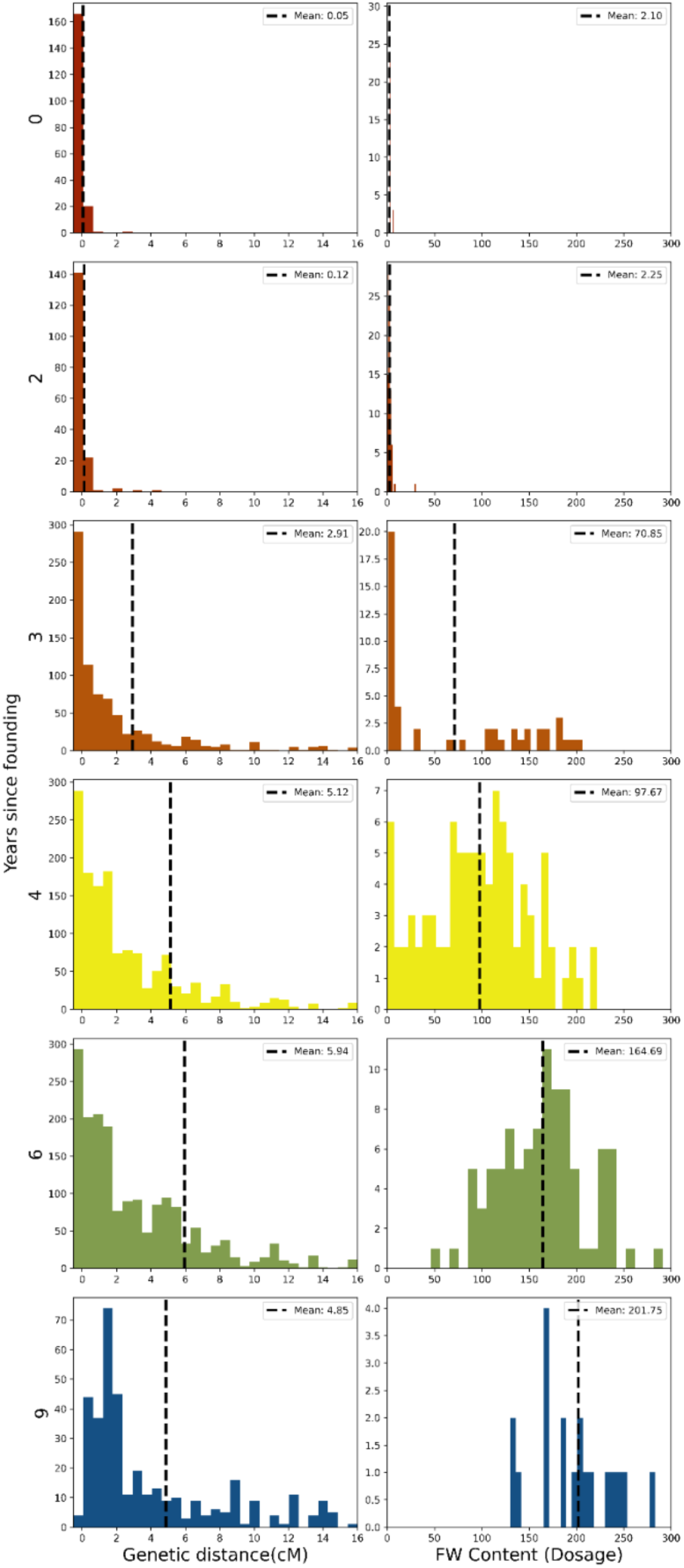
Distance between freshwater adaptive loci and freshwater content at each locus during early stages of freshwater adaptation: **(A)** Distribution of measured distance between first SNP of the first locus and last SNP of the last locus of a contiguous block. A contiguous block is defined as two or more consecutive loci with freshwater adaptive alleles **(B)** Freshwater content is estimated as the proportion of freshwater dosage all loci in a time point. A locus with homozygous freshwater has a dosage of 2, heterozygote has 1 and marine homozygote has 0. The black dashed line is the mean in both A and B. FW = Freshwater

In stark contrast to the SC2013 sample, in the third year after founding (SC2014, F2 generation, fig. 1D), nearly half (21 out of 47) of the individuals in our sample possessed large numbers of freshwater-adaptive alleles (fig. 1F, 2B, S2). The mean proportion of freshwater-adaptive alleles in SC2014 was significantly different from that of SC2013 (mean proportion of freshwater alleles in SC2014=0.1222 vs SC2013=0.0048; *t* = 6.3156, df = 47.03, *p* = 8.923e-08; fig. 2B, S2). The distribution of the number of freshwater-adaptive alleles per individual in the SC2014 sample was bimodal (fig. S2). Therefore, to aid our understanding of genomic change during adaptation to freshwater, we used this bimodal distribution to categorize individuals as *jackpot carriers* when they had more than 10% of the freshwater-adaptive alleles and *non-jackpot individuals* if they had fewer. Based on this classification, there were no jackpot carriers in the RS2019 and SC2013 samples (i.e., 0%), while there were 21 out of 47 in SC2014 (i.e., 45%), 78 out of 96 in SC2015 (i.e., 81%), 95 out of 96 (i.e., 95%) in SC2017, and all 20 in SC2020 (i.e., 100%, Table S1). We observed that 27 (21 jackpot carriers and six non-jackpot individuals) out of the 47 individuals in SC2014 had contiguous blocks with freshwater-adaptive alleles. These contiguous blocks were mostly heterozygous (fig. S4) and spanned 0.07Mb[0.044cM] to 22.33Mb [70.48cM], with a mean of 2.40Mb[2.9cM] (fig. 2A, S3, Supplementary table 2).

Threespine stickleback in the Cook Inlet lakes become reproductively mature at around one or two years old,^27–30^ and results from Kurz et al.^24^ indicated that by 2014, the captured individuals were F2 (fig. 1E). Therefore, the observed jackpot carriers cannot be a result of re-assembly via recombination from anadromous founders with only a few freshwater-adaptive alleles. Jackpot carriers must have been present in the founding population at such low frequencies that they were undetectable in our limited RS2019 or SC2013 samples. Based on previous estimates of the frequency of jackpot carriers in ancestral marine environments of 0.1%^12,17^, the probability of including at least one jackpot carrier in a sample of 96 individuals is only 9% (assuming binomial sampling probabilities). The rapid increase in the frequency of jackpot carriers from undetectable in the F1 generation (i.e., SC2013), which was born and spent its entire life in the lake, to almost 45% in the F2 generation (i.e., SC2014) was likely a result of their markedly greater fitness in the freshwater environment compared to individuals with few freshwater-adaptive alleles after founding.

Low survival and/or low reproductive success of the vast majority of non-jackpot carriers during years two and three after founding resulted in a population bottleneck that was manifested by a markedly reduced catch per unit effort (CPUE), a relative measure of abundance (fig. 3A). In the SC2013 sample, the decline in CPUE was consistent with a decrease in genome-wide genetic diversity compared to RS2019 sample (Watterson’s Θ=0.0058 vs 0.0047 for SC2013) (fig. 3B). CPUE further declined in SC2014, and a correspondingly more significant reduction in genetic diversity of almost 50% compared to RS2019 (Θ = 0.0030 for SC2014) (fig. 3A, 3B). This population decline in Scout Lake caused Tajima’s D to become increasingly positive in the SC2013 (D=0.263) and the SC2014 (D=1.618) sample despite the RS2019 sample having a negative value (D=-0.722) that likely reflects a population expansion in the ancestral environment. The genome-wide site frequency spectrum (SFS), estimated using ANGSD^31^, indicates progressive, genome-wide loss of singletons, with a particularly drastic distortion of the SFS in SC2014 for rare alleles, and is consistent with the dramatic population decline within the Scout Lake population (fig. 3C).

**Fig. 3:**
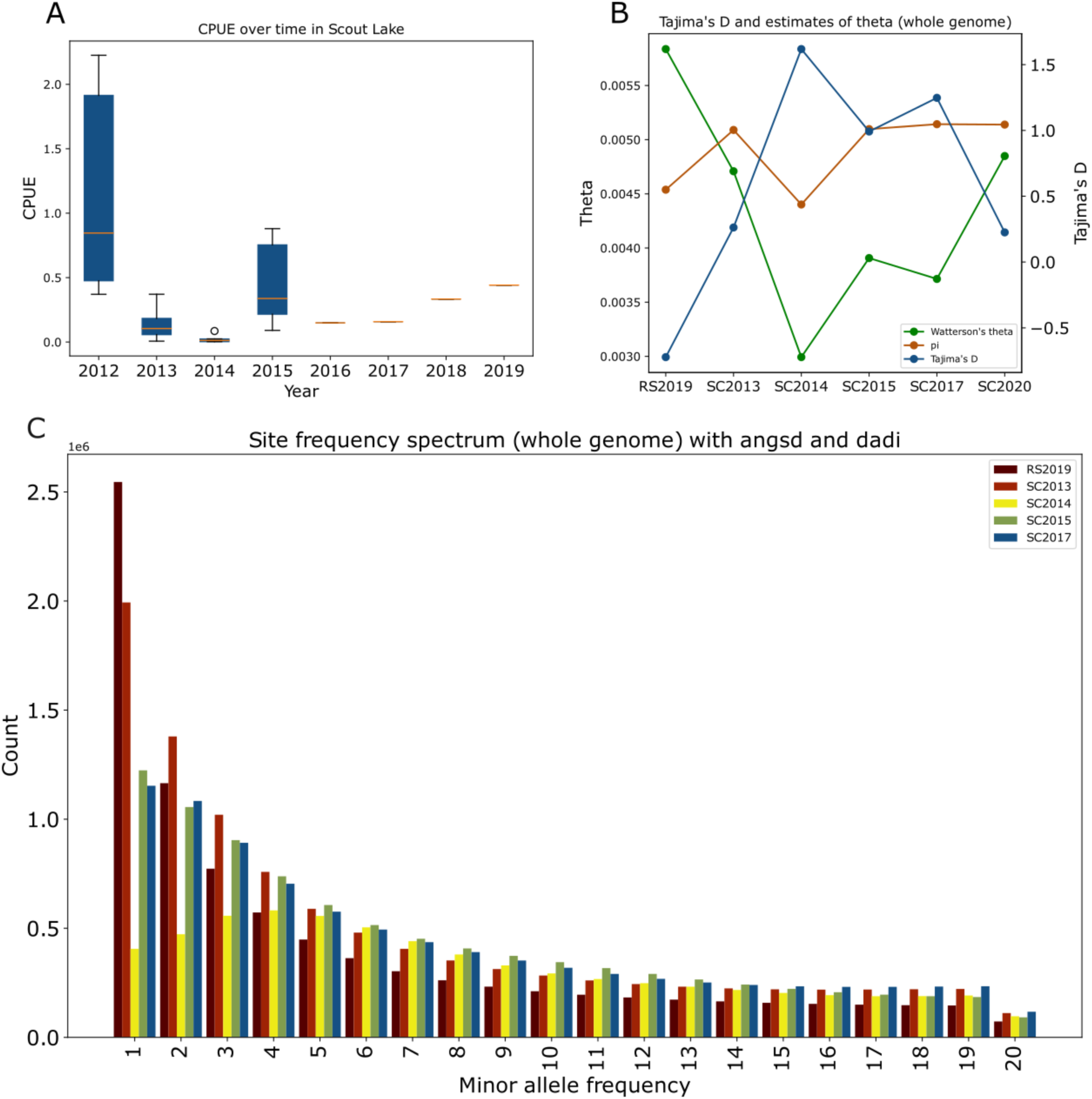
Measures of demographic changes during rapid adaptation in Scout Lake: **(A)** Catch per unit effort (CPUE). CPUE was measured as the number of stickleback caught per trap hour. **(B)** Summary statistics (Tajima’s D, theta and nucleotide diversity) calculated from the whole genome for Rabbit Slough, SC2013, SC2014, SC2015, SC2017 and SC2020 **(C)** Site Frequency spectrum (SFS) estimated from the whole genome. The SFS was folded in dadi and projected down to 40. For clarity, we only show the first 20 polymorphic sites in the SFS. We excluded SC2020 from the SFS in C because of the differences in depth of coverage with the other samples. The SFS with the SC2020 sample can be found in fig. S12 and the unprojected SFS with all polymorphic sites can be found in fig. S13.

After this bottleneck, the CPUE increased, and there was a relative increase in the frequency of rarer variants in the SFS for the SC2015 and SC2017 samples compared to SC2014. The number of freshwater-adaptive alleles increased consistently over time in samples collected after the bottleneck; the mean proportion of freshwater-adaptive alleles in SC2015, SC2017 and SC2020, were 0.2078, 0.3504, 0.4292, respectively (fig. 1F, 2B, S2). However, the size of freshwater haploblocks, as measured by the number of loci per block, or the haploblock length in Mb or cM (fig. 2A, S3), did not change in concert, as would have been expected if recombination had been bringing freshwater-adaptive alleles together from multiple parents each generation to form increasingly larger haploblocks. For instance, the mean haploblock length of contiguous loci with freshwater-adaptive alleles was 2.67 Mb [(range: 3e-05Mb-23.97Mb) in SC2015, 2.93 Mb in SC2017, and 2.69 in SC2020. Yet, the homozygosity of freshwater-adaptive loci increased over time from SC2014 (fig. S4), which could reflect the mating between largely heterozygous individuals. These results demonstrate that haploblocks of freshwater-adaptive alleles were not re-assembled through progressive recombination over many generations, but instead that the frequencies of freshwater-adaptive alleles increased after the F1 generation by matings predominantly between the descendants of jackpot carriers, which were present in the founders.

### Biological kinship networks and inbreeding during freshwater adaptation

Our sequence data from multiple whole genomes during the earlier years after the Scout Lake population was founded provided a unique opportunity to explore the genealogical relationships among individuals as the population adapted to the new freshwater environment and to infer the dynamics of haploblock inheritance. We, therefore, estimated pairwise biological relatedness among our low-coverage genomes using READv2^32,33^ and ngsRelate^34^. The estimates of relatedness coefficients from the two methods were highly correlated (r>0.92 for all time points; fig. S5). However, in situations in which there is increased allelic sharing compared to expectations from panmixia due to inbreeding, we might expect READv2 relatedness coefficients to be more robust as the approach relies on observed rates of allele mismatching, while ngsRelate utilizes allele frequencies under the assumption of Hardy-Weinberg equilibrium, leading to potentially inflated values of relatedness (Methods, Supplementary Section 8). Therefore, we report the results based on READv2, which identifies up to third-degree relatives. First-degree relatives can either be parents-offspring pairs or siblings. Second-degree relatives include grandparent-grandchild pairs and avuncular pairs. Third-degree relatives include first cousins, great aunts/uncles, grandniece/nephews, and great-grandparents. We can distinguish the two types of first-degree relatives but not the various types of second or third-degree relatives (Supplementary Section 8).

We also estimated individual inbreeding coefficients, *F*, with ngsRelate, which can estimate inbreeding from low-coverage genomes (up to approximately 4X^34^). A down-sampling experiment of the high-coverage genomes from the SC2020 sample demonstrated that ngsRelate underestimates inbreeding coefficients from low-coverage of 1X by a factor of ∼50% compared to high-coverage (∼30X) whole genome data (Supplementary Section 9, fig. S8); thus the reported inbreeding coefficients from the low-coverage genomes from the SC2013 to SC2017 samples are likely underestimated. All pairwise relationships detected using READv2 and NGSrelate are provided in Supplementary table 5).

There were no individuals with up to third-degree relatedness in the RS2019 sample, suggesting that the founders of the Scout Lake population were drawn from an outbred anadromous population with a large effective size (N_e_). We collected the RS2019 sample in 2019, and it is not a subset of the individuals used to establish the new Scout Lake population in 2011, although we sampled the fish similarly. Despite possessing identical proportions of freshwater-adaptive alleles as RS2019 (fig. 1F), the SC2013 sample had two first-, 38 second- and five third-degree relatives (fig. 4A, S6A), reflecting that this population is descended from a small number of fish with a much more limited choice of mates than the ancestral anadromous population. Both first-degree relatives in the SC2013 sample were siblings, which aligns with our generation inference based on phenotypic data (fig. 1E) that showed that it is unlikely that one-year-old F1s reproduced in 2012.

**Fig. 4:**
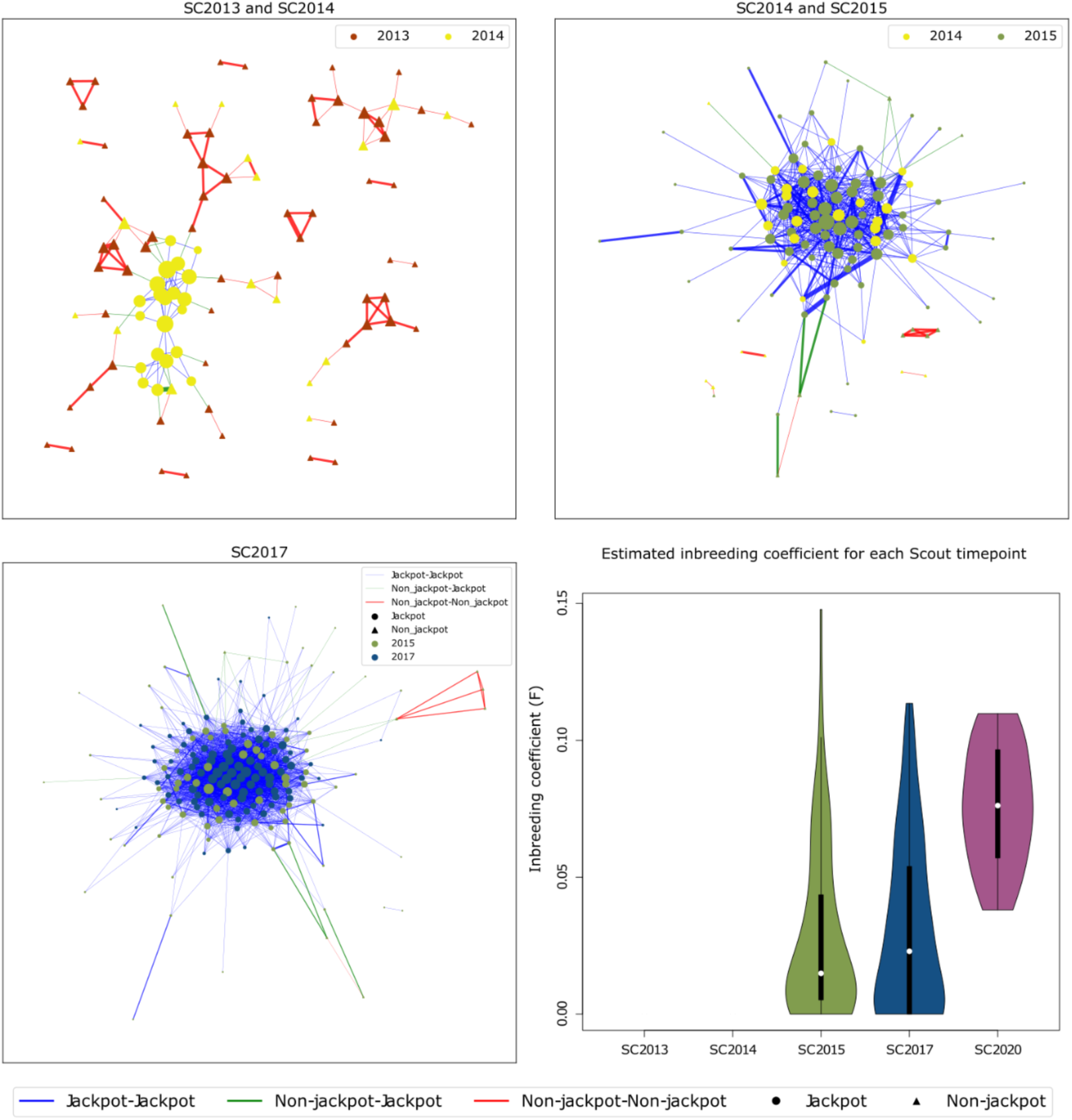
Inter-year relationships and inbreeding: **(A)** Relatedness between SC2013 (2 years since founding) and SC2014 (3 years since founding). **(B)** Relatedness between SC2014 (3 years since founding) and SC2015 (4 years since founding) **(C)** Relatedness between samples SC2015 ( 4 years since founding) and SC2017 (6 years since founding) **(D)** Inbreeding coefficients estimated from ngsRelate at the various time points. No inbred individual was observed in SC2013 and SC2014, 2 and 3 years after founding. In panels A, B and C, the thickness of edge reflects the degree of relatedness between the nodes: the thickest edge first-degree, intermediate second-degree and the thinnest edge third-degree relatives. The size of node increases with increasing number of relatives.

Some related pairs involved one individual from the SC2013 sample and the other from the SC2014 sample. There were 36 third-degree and two second-degree relative such pairs (fig. 4A). Thirteen out of the 36 third-degree pairs were between a jackpot carrier from SC2014 and non-jackpot carriers in SC2013, suggesting that jackpot carriers mated with non-jackpot individuals early on.

There were one first-, one second-, and 58 third-degree relatives in the SC2014 sample. While the majority of relatedness estimates within the SC2013 sample (fig. S6A) formed small networks, each containing only a few individuals, out of a total of 60 related pairs within 2014, 57 were connected within a single large network (fig. S6b). The one first-degree relationship was a sibling pair. Apart from four relationships between jackpot and non-jackpot individuals, this network’s remaining pairs were jackpot carriers (fig. 4A, S2B). All jackpot carriers (21 out of 47 individuals) identified in the SC2014 sample were connected within this single network. The third-degree relationships of jackpot carriers sampled in 2014 are likely to be cousins or half-avuncular since the typical lifespan of Alaskan Threespine Stickleback is one to two years^29^.

We observed one first-degree (sibling pair), nine second-degree, and 269 third-degree pairs of relatives between specimens from the SC2014 and SC2015 samples. All except two third-degree relationships were in a network that consisted of jackpot carriers (all the jackpot carriers in the network observed in SC2014, and 57 out of 78 jackpot carriers in SC2015) (fig. 4B). Within the SC2015 sample (fig. S6C), there were one first-degree (sibling pair), 36 second- and 298 third-degree pairs. These 335 connections were between 77 out of 96 sampled individuals in SC2015, 55 of which were jackpot carriers (there were 78 jackpot carriers in SC2015). Unlike in the SC2013 and SC2014 samples, we found that some individuals in SC2015 showed evidence of inbreeding (fig. 4D), with *F* values ranging from 0.0 to 0.149 (mean=0.0289). Jackpot carriers were more inbred (mean=0.032, range=[0.0, 0.149]) than non-jackpot individuals (mean=0.0134, range=[0.0001, 0.0362]), [t=4.08, *p-value*= 0.0001]. The jackpot carriers observed in SC2015 were mostly related to the SC2014 jackpot carriers, suggesting that the SC2015 jackpot carriers descended from the network of jackpot carriers in SC2014.

The individual with notable contiguous blocks of freshwater-adaptive alleles in the SC2013 sample was a third-degree relative of an individual in the SC2014 sample with a similar proportion of freshwater alleles. No relatives from this pair were found in the SC2015 sample or later, and this lineage presumably died out in 2014 or failed to proliferate enough to be represented in subsequent samples, possibly because they possessed too few freshwater-adaptive alleles. Thus, their freshwater content of ∼7% was insufficient to contribute to the growing population from 2014 and beyond, justifying our criteria of defining a jackpot carrier as individuals with ≥ 10% freshwater-adaptive alleles.

The network of relationships between the SC2015 and SC2017 sample provided further evidence that population growth was driven by jackpot carriers (fig. 3C), with 34 second- and 1201 third-degree pairs in 165 out of 192 (i.e.. 85.9%) individuals from SC2015 and SC2017. All relationships, except seven third-degree and one second-degree relative, were between jackpot carriers. By 2017, only one out of 96 sampled individuals was not a jackpot carrier, and 93 out of the 95 jackpot carriers were in a single network (fig. S6D). The individuals sampled in SC2017 had an average inbreeding coefficient of 0.0316 (range [0.0, 0.1121]). None of the 20 individuals in the SC2020 sample were first-, second- or third-degree relatives, though we identified three third-degree relatives between SC2017 and SC2020 individuals (Supplementary table 5). To test if this lack of relatedness within SC2020 was significantly lower than in SC2017 or simply a result of the smaller sample size (SC2017 n = 96, vs. SC2020 n=20), we performed a permutation test by randomly sampling 20 individuals from the SC2017 sample 1000 times and observing the number of pairwise relatives of at least third-degree. In no permutation did we observe 20 random individuals from SC2017 with no relatedness (mean=57.2, min=16, max=112, fig. S7). This result suggests that the lack of relatedness in the SC2020 sample was likely to be a result of population growth that reduced the probability of sampling relatives and not a result of the small sample size. Despite the deficiency of related pairs, individuals in SC2020 were, on average, 0.0758 inbred (range [0.038, 0.1098]). The dearth of relatives within SC2020 suggests that the population had become large enough to allow more mate choice by this time, with the increased *F* reflecting the legacy of inbreeding from the preceding generations.

Our kinship analysis demonstrated that jackpot carriers mated with non-jackpot individuals early on in the adaptive process, prior to the bottleneck in 2014. In order to examine the extent to which founding non-jackpot individuals contributed freshwater-adaptive alleles to the later post-bottleneck population, we identified loci where all 21 jackpot carriers found in the SC2014 sample were homozygous for the oceanic allele. There were four such unlinked loci. These loci included an 11Kb locus on chromosome IV, a 2Kb locus on chrIX, and 11Kb and 19Kb loci on chrXIX. We then determined if any individual in the subsequent timepoints (SC2015, SC2017 and SC2020) had inherited freshwater alleles at these loci. We found only five heterozygous individuals across all four loci in all the subsequent samples (N=212). However, there were no freshwater adaptive alleles in the non-jackpot individuals observed in the SC2013 and SC2014 samples at these four loci. This indicates that non-jackpot individuals within the founders contributed minimally to the pool of adaptive alleles circulating after the bottleneck.

### Rapid freshwater adaptation was not sex-specific

The closely related network of jackpot carriers observed in the SC2014 through SC2017 samples indicated they descended from a few male or female ancestors in the founding population. If this is true, we expect that the observed jackpot carriers in SC2014 and their subsequent offspring would be enriched with the mitochondrial (mtDNA) or Y-chromosome haplotype(s) carried by their common ancestor(s). Filtering for polymorphic sites on the mtDNA with frequency >5% across all genomes, we identified nine high-confidence haplotypes, hereafter haplotypes A through I. Given the low-coverage of our genomic data, we could not call haploid genotypes on the non-recombining portion of the Y chromosome (NRY) with high confidence (mean coverage is expected to be half the autosomes), with substantial missing data across individuals for any particular site. We, therefore, developed an imputation approach based on the complete linkage of the NRY (Supplementary Section 10) and filtering for regions of the NRY showing consistent coverage, similar to an approach adopted in Lippold et. al^35^. This approach allowed us to identify four high-confidence NRY haplotypes defined by three polymorphic sites.

The E haplotype of the mtDNA was found at low frequencies (<10%) in the RS2019 and SC2013 samples, and the GGC NRY haplotype was absent at these timepoints. However, these two haplotypes rapidly increased in frequency from 2014 onwards to become the highest frequency haplotypes in 2017 (60% and 38%, respectively) (fig. 5). We performed an Exact Test of Population Differentiation under the null hypothesis that there is no difference between the sampled timepoints based on haplotype frequencies^36^. This test demonstrated that at a significance level of 0.05, SC2013 was not differentiated from RS2019 for either mtDNA or NRY (*p-value*=0.24741 for mtDNA and 0.12536 for NRY, Supplementary table 3). However, for mtDNA, the SC2014, SC2015 and SC2017 samples were significantly differentiated from the SC2013 sample (*p*<0.05). For the NRY, the SC2015 and SC2017 samples, but not the SC2014 sample, were differentiated from the SC2013 sample (Supplementary Table 3). These results point to significant shifts in haplotype frequencies at both mtDNA and NRY over time.

**Fig. 5:**
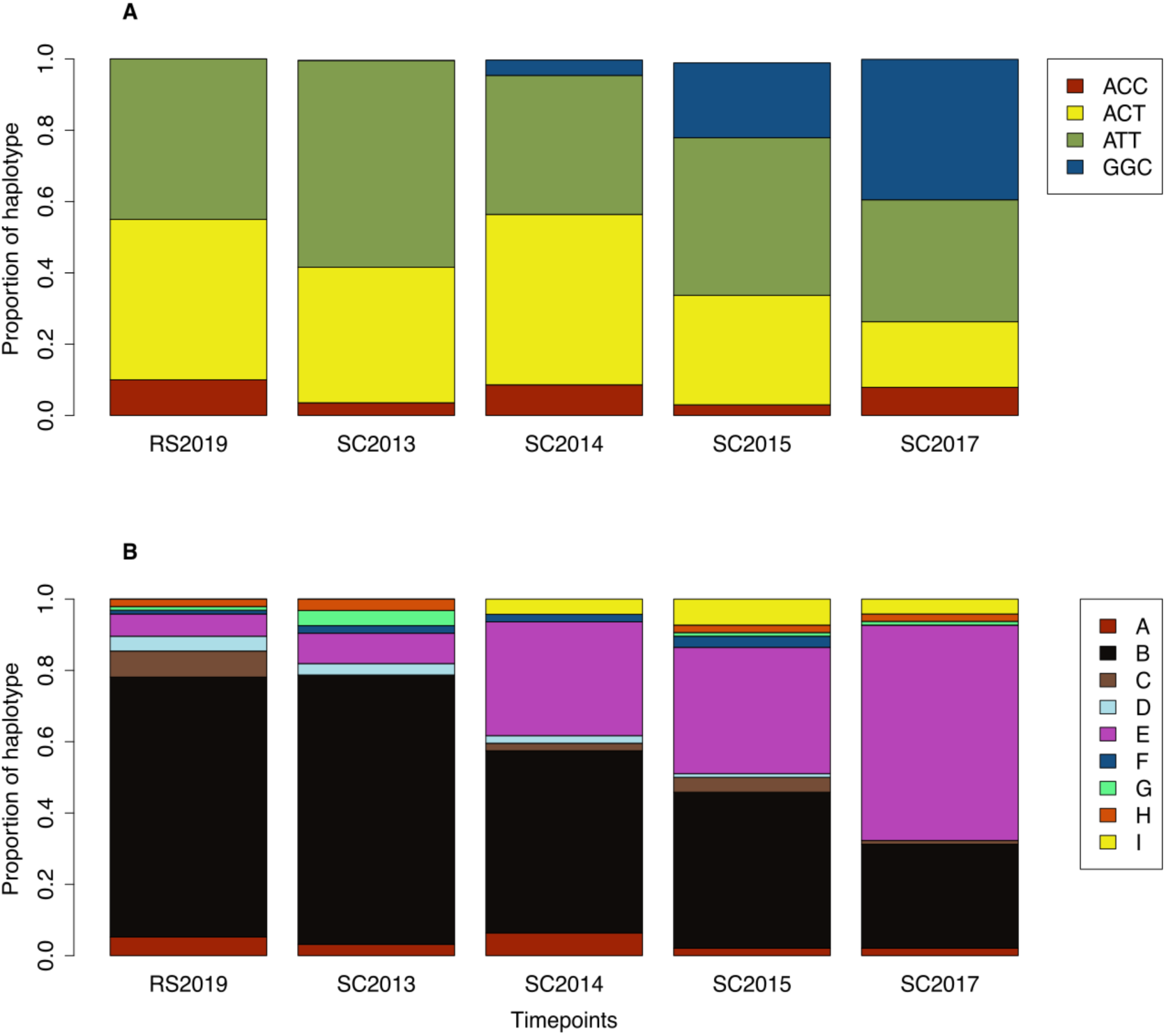
Haplotype diversity Y-chromosome (NRY) and mitochondrial DNA (mtDNA). **(A)** Proportions of NRY haplotypes (ACC, ACT, ATT and GCC) in the sampled. GCC haplotype only in 2014 (3 years since founding). **(B)** Proportions of haplotypes (A to I) in the sample. We excluded SC2020 from this analyses due to small sample size.

Twenty-five out of 26 male individuals possessing the GGC haplotype on the NRY and 101 out of 107 male and female individuals with the mtDNA E haplotype were jackpot carriers. Among jackpot carriers, 51% possessed the E mtDNA haplotype compared to 8.5% in non-jackpot individuals (fig. S9). The frequency of the GGC NRY haplotype in jackpot carriers was 29% (25 out of 86 individuals) and 0.8% (1 out of 128 individuals) in non-jackpot individuals. Thus, in both mtDNA and NRY loci, specific haplotypes were common in jackpot carriers compared to non-jackpot individuals, and increased progressively over time. These observations are consistent with selection being driven by both male and female jackpot carriers rather than a sex-specific process.

### Patterns of polymorphism, selection and genetic load during adaptation to freshwater

When a population experiences a decline in size, the increased probability of inbreeding can increase genetic load^37–39^ and reduce mean population fitness^39,40^. Rare, deleterious, recessive alleles previously masked in heterozygotes become exposed as homozygotes and reduce individual fitness^37,38,41^. To determine how the dynamics of the observed bottleneck affected the genetic load of the Scout Lake population, we estimated the SFS genome-wide as well as at 4-fold (silent sites) and 0-fold (amino acid replacement) degenerate sites for samples from the time series (Supplementary Section 11).

As expected, 0-fold sites generally demonstrate approximately a third of the diversity of 4-fold and genome-wide sites at all samples, presumably due to the effects of purifying selection (Fig. 6F, S10). RS2019 showed a slight excess of singletons at 0-fold sites compared to 4-fold and genome-wide sites, plausibly due to the segregation of slightly deleterious alleles in the ancestral anadromous population (Fig. 6A, S10). During the bottleneck from 2013 to 2014 (SC2013 and SC2014 samples), there was a similar reduction in Watterson’s Θ at 0-fold and 4-fold sites and genome-wide (30%, 34% and 25%, respectively), indicating that a demographic factor, the population bottleneck, shaped most of the pattern of diversity during this time (fig. 6F).

**Fig 6:**
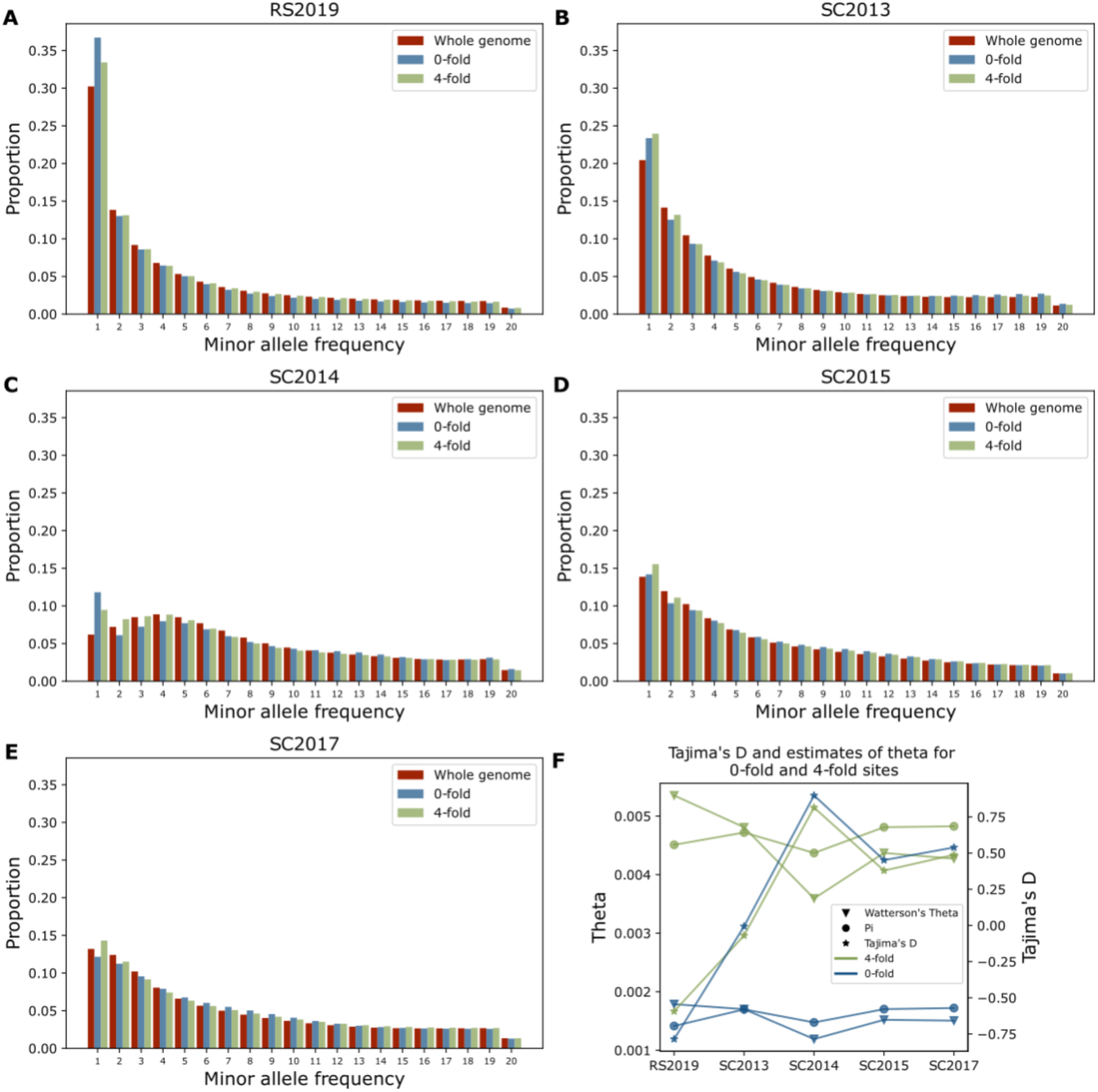
**(A-E)** Site Frequency Spectrum (SFS) at the whole genome, 4-fold and 0-fold degenerate sites at various time points. **(F)** Estimates of theta and Tajima’s D for the various time points at 4-fold and 0-fold sites.

However, within the SC2014 sample a notable increase in singletons at 0-fold sites relative to that expected at genome-wide sites was detected, with 4-fold sites being intermediate, presumably reflecting linkage to 0-fold sites (Fig. 6C). Compared to non-jackpot individuals, the SFS of jackpot carriers was skewed towards more common variants with no significant increase in 0-fold singletons, likely reflecting the close genealogical relationships between these individuals (fig. S11). The singletons in non-jackpot carriers reflected the excess of singletons at 0-fold sites in the adult F2 generation (SC2014). We could, therefore, attribute the observed singletons at 0-fold sites almost entirely to non-jackpot carriers. However, this excess of singletons at 0-fold sites was not observed in the samples collected from 2015 and 2017 (SC2015 to SC2017). As described above, in SC2015 and SC2017, we also observed high levels of relatedness and inbreeding as well as increased homozygosity (fig. S4). Thus, slightly deleterious 0-fold sites were likely being purged from the population as a result of a) non-jackpot individuals failing to leave descendants except when they mated with jackpot carriers and b) an increased chance of recessive mutations being homozygous and thus being selected against.

### Phenotypes of jackpot carriers and non-jackpot individuals in SC2014

Some phenotypic divergence between newly founded freshwater Threespine Stickleback populations and their oceanic ancestors are conspicuous^15,29^. One generation after freshwater colonization (i.e., SC2013), the female standard lengths (SL) in Scout Lake declined significantly^24^, although SL is phenotypically plastic^42^. Other traits such as lateral plate morph, dorsal fin ray number, and gill raker number are highly heritable^15,43,44^. We compared the standard length, lateral plate morph and dorsal fin ray number between jackpot and non-jackpot individuals in the SC2014 sample and found no significant phenotypic differences between them (fig. S14). There were 16 out of 21 jackpot carriers that possessed a freshwater allele at the locus that contains the *EDA* gene, which strongly influences the lateral plate morph in Threespine Stickleback^10,45,46^ compared to one out of 26 non-jackpot individuals (X^2^ (1, N=47) =26.85, *p-value* = 2.197e-07). However, the freshwater alleles in these 16 jackpot carriers were all in a heterozygous state. Thus, they would not be expressed if the allele is recessive for all phenotypes that it influences^47^, and all but one fish (partially plated) had a high plate morph phenotype.

Physiological phenotypes such as fatty acid metabolism may be crucial during adaptation to freshwater environments by oceanic Threespine Stickleback^48,49^. Ishikawa et al.^49^ found evidence suggesting that increased copy number variants for the fatty acid desaturase (*fads2*) gene play a key role in colonizing freshwater environments by oceanic Threespine Stickleback. We tested the hypothesis that the Scout Lake population had elevated copies of *fads2* than the ancestral anadromous ancestors using our high-coverage SC2020 sample (Methods). We calculated the average coverage across individuals at all the autosomes, the sex chromosome (chromosome XIX:12500000-20618466) and two of the three known copies of the fads2 gene (chromosome XII:13777552-13780899 and chromosome XIX:17503349-17507332). We summed the average coverage of the *fads2* gene copies on chromosome XII and XIX together. We also summed the average coverage on the autosomes and chromosome XIX. We then computed the ratio of the two quantities, with the expectation that a higher copy number of *fads2* will produce an elevated ratio more than 1. The ratio of the two quantities was 0.87. There were two males with an extra copy of *fads2* on their chromosome XIX (supplementary table 8). We also estimated *fads2* copy number of 20 high-coverage genomes from a sample from Rabbit Slough collected in 2009^12^. The ratio of the two quantities in this sample was 0.80. The *fads2* copy number in the specimens in the Rabbit slough 2009 sample and SC2020 sample were not significantly different for either the *fads2* copy on chromosome XII (ANOVA, F_1,38_ = 2.2, P = 0.14) or chromosome XIX (ANOVA, F_1,38_ = 0.92, P = 0.34). These results suggest that *fads2* copy number does not significantly increase during the first few generations of freshwater adaptation in Scout Lake population.

## DISCUSSION

### Jackpot carriers mediate rapid freshwater adaptation

We studied the rapid adaptation of an anadromous Threespine Stickleback population to a new freshwater environment by comparing genomic variation during the first few generations after the population was founded and in contrast to the population from which it was derived^21^. Adaptation appears to have been driven primarily by a few individuals with large haploblocks of freshwater adaptive alleles (i.e., jackpot carriers) that produced families that become predominant in the population during the first nine years after founding. Jackpot carriers were presumably present among the anadromous Rabbit Slough founders released into Scout Lake. However, they were at such low frequencies that they were undetected in a representative sample of the ancestral Rabbit Slough population (i.e., RS2019) and the sample from the F1 generation in the lake (i.e., SC2013). These jackpot carriers had markedly higher fitness in Scout Lake compared to non-jackpot individuals, driving the process of freshwater adaptation via population growth from just a few ancestors. This process conflicts with the transporter hypothesis^16^ in that freshwater adaptation occurs via the spread of large pre-existing haploblocks of freshwater-adaptive alleles within a few generations in the lake rather than during many generations of recombination of freshwater-adaptive alleles derived from numerous founders with a few freshwater-adaptive alleles each.

The sudden appearance of jackpot individuals in 2014 could be due to a) F2 jackpot carriers having an increased rate of survival throughout their life-cycle compared to non-jackpot individuals or b) F1 jackpot carriers having increased reproductive success. Previous experimental work has demonstrated that in freshwater families (derived from both marine and freshwater parental ancestries) which were completely plated grew slower than those families with reduced plates^50^. Although growth rate potentially impacts survival over the winter period^51^, we found no size difference between jackpot carriers and non-jackpot individuals from the 2014 sample. On the other hand, there appears to be no difference in hatching rates between marine and freshwater families in freshwater^50^, which would be expected given anadromous fish by definition breed in freshwater. Thus, there is little evidence that would support either hypothesis of greater survivorship or greater reproductive success of the jackpot F1s. Additional research is therefore required to discriminate between these two alternatives.

If the jackpot carriers observed in the SC2014 sample descended from anadromous-freshwater hybrids that we released into the lake in 2011, we expect approximately 50% of their alleles at freshwater-adaptive loci to be freshwater if they were first^-^generation hybrids, 25% for first-generation backcrosses to anadromous stickleback, and 12.5% for second-generation backcrosses, assuming that rare hybrids subsequently backcrossed to non-jackpot carriers with anadromous ancestry that are likely typical of Rabbit Slough fish. The content of freshwater alleles in the jackpot carriers identified in the SC2014 sample ranged from 12.1% to 49.2%, with a mean of 33%, suggesting that these individuals were first to third-generation anadromous-freshwater hybrids. The presence of recent generations of hybrids in the founding population indicates significant gene flow from a freshwater population to the anadromous Rabbit Slough population that we used to found the Scout Lake population, and this is probably a key element of the process by which oceanic Threespine Stickleback adapt to new freshwater environments, as suggested by the transporter hypothesis^10,16^.

An alternative scenario for the presence of recent hybrids between anadromous and freshwater stickleback in the Scout Lake samples is that the jackpot carriers were produced by matings between anadromous founders and very rare freshwater individuals from the original Scout Lake population that had survived the rotenone treatment of the lake. We tested this hypothesis by creating hybrids from established freshwater populations from the Pacific Coast and Rabbit Slough individuals (see Supplementary Section 13). The distribution of freshwater-adaptive alleles in these hybrids was 50 to 63% compared to the 12.5% to 49.5% content we found in the individuals we sampled in the SC2014 sample, suggesting that, the jackpot carriers observed in SC2014 sample were probably recent oceanic-freshwater hybrids, or their progeny. These were likely to have been present among the Rabbit Slough founders and not hybrids between survivors from the original freshwater Scout Lake population and released anadromous individuals. This latter scenario is further supported by previous studies that have found jackpot carriers in samples collected from marine environments^12,17^.

Bassham *et al.*^17^ studied multiple natural freshwater Threespine Stickleback populations that were more than 50 years old^52^ and suggested that freshwater-adaptive alleles could enter new environments as colocalized haplotypes (i.e., as large haploblocks) but that multiple colonization events would be necessary to reassemble fully freshwater ecotypes. However, our results show that introducing about 3000 anadromous sticklebacks within 30 days was sufficient to include enough jackpot carriers for adaptation to proceed. The introduction of about 3000 anadromous sticklebacks from Rabbit Slough to two other Cook Inlet lakes in 2009^53^ and 2019^22^ also established rapidly adapting populations. Assuming a frequency of jackpot carriers of 0.1% in the oceanic environment, as estimated previously^12,17^, the probability of observing at least one jackpot carrier under binomial sampling in the approximately 3000 anadromous fish we transplanted into Scout Lake is 0.95. However, the probability of capturing at least 5, 7 or 10 jackpot carriers is only ∼0.2, 0.03 and 0.001, respectively. Given that the jackpot carriers in the Scout Lake population in 2014 and subsequent generations in 2015, 2017 and 2020 primarily consist of a closed network of related jackpot carriers, it appears that adaptation of anadromous stickleback to freshwater can occur with approximately five descendants of recent oceanic-freshwater hybrids in the founding anadromous populations, whether spread over multiple small colonization events, as proposed previously^52^ or in a single event, as observed in Cheney and Scout lakes^15^.

Non-jackpot individuals may also be required to successfully colonize freshwater by anadromous stickleback. At such low frequencies, the probability of jackpot individuals mating with each other during the earliest years may be low, particularly when there are no noticeable phenotypic differences between jackpot and non-jackpot individuals (though behavioral differences may increase the chance of this). Mating occurred between jackpot and non-jackpot individuals before 2014, as we found relatedness compatible with half-sibling and half-avuncular relationships in both 2013 and 2014. Mating with multiple non-jackpot individuals would provide a mechanism for jackpot haploblocks to increase rapidly in frequency in subsequent generations, especially given that Threespine Stickleback nests can contain more than 300 eggs^54,55^. Low recombination rates at freshwater-adaptive loci ^12,19^ will also ensure that large heterozygote haploblocks are inherited largely intact by numerous offspring during each generation. Thus, jackpot haploblocks could increase their frequency by orders of magnitude in just one generation of mating simply by jackpot carriers pairing with non-jackpot individuals.

### The consequences of bottlenecks on genetic load during rapid adaptation

Previous studies have proposed that population bottlenecks may be important during freshwater adaptation by oceanic Threespine Stickleback^56,57^. These bottlenecks have been inferred from observed reduction in genetic diversity in established freshwater populations^56–58^. Here, we present the first direct observations of the temporal dynamics of stickleback adaptation to freshwater during the first nine years. This process in Scout Lake involved a bottleneck that occurred during the third and fourth year after founding, likely as a result of reduced fitness of non-jackpot individuals that made up the majority of the founding population. The population then recovered as a result of higher fitness of a few jackpot carriers.

The resulting reduction in population size during the early years after founding coincided with a major shift in the genetic composition of the individuals that remained in the lake, indicating the population decline was non-random. However, such bottlenecks also increase the probability that a population will lose the genetic variation required to adapt, leading to extinction. Population growth from a few related individuals can decrease genetic variation and increase the genetic load of a population. The genetic load could either be expressed phenotypically or masked^38^ in diploid populations, and bottlenecks can increase phenotypic expression of a population’s genetic load^37–39^ by exposing rare, recessive deleterious alleles masked in heterozygotes.

Inbreeding after the bottleneck could increase the frequency of rare deleterious alleles and their probability of homozygosity^37,38,41^. Such a scenario can lead to one of two outcomes: a) the population could be at risk of extinction due to the increase in the fixation probability of recessive deleterious mutations^59–61^ through drift, or b) negative selection could be more effective at eliminating slightly deleterious mutations, as selection is more potent against recessive or partially recessive mutants in the homozygous state^62^.

Our analysis of 0-fold and 4-fold degenerate sites as a proxy for genetic load suggests that inbreeding after the bottleneck may have facilitated the removal of slightly deleterious mutations by unmasking them as homozygotes. While increased homozygosity can put the population at risk of extinction, the explosive growth associated with a strong positive selection of jackpot carriers may have balanced that adverse effect and increased the chances for the population to survive the bottleneck. This process could, therefore, increase the population’s overall adaptability by removing deleterious variants that circulated in the anadromous ancestor alongside the positive selection of freshwater alleles, at least during the early stages of the adaptive process, before new deleterious mutations emerge.

Ernst Mayr emphasized the influence of demographic bottlenecks and resulting genetic drift (founder effect) on phenotypic divergence and speciation^63,64^. In addition Eldredge and Gould incorporated the founder effect in their punctuated equilibrium model^65^, while a bottleneck was central to Carson’s flush-crash modeling of speciation by Hawaiian *Drosophila* species^66^. In the invasive blowfly, *Calliphora vicina,* there is evidence that bottlenecks could underlie their expansion into new environments^67^. Bottlenecks may also play crucial roles in divergence and speciation in other vertebrates, such as the *Cyprinodon tularosa*^68^ and in plants such as *Picea abies*^69^. These findings in diverse organisms indicate that bottlenecks influence adaptation across the tree of life and may not be peculiar to rapid adaptation to freshwater by anadromous Threespine Stickleback.

### Physiological but not morphological features may be necessary during the early stages of freshwater adaptation

Despite our observation that nearly half of the individuals within the SC2014 sample were jackpot carriers, we observed no significant differences in morphological phenotypes between jackpot carriers and non-jackpot individuals for selected traits that differ characteristically between oceanic and freshwater resident populations. This result suggests that physiological traits that underlie osmoregulation and metabolism may be more critical during the earliest stages of freshwater adaptation.

Even increased frequency of low armor plate morphs at the expense of ancestral complete morphs, a consistent, major signature of the classic freshwater ecotype^50,70^, did not occur in the Scout Lake population during and shortly after the population bottleneck, despite likely normal predation pressure in the lake. (Scout Lake was re-stocked with invertebrates, and there are avian predators.) Morphological phenotypes divergent between freshwater and oceanic ecotypes may respond more slowly to directional selection on morphological traits or exert a more significant fitness effect at later stages of adaptation. This situation may explain why traits like loss of armor plating are recessive and likely become expressed after the alleles increase in frequency and are then expressed when individuals inherit two of these recessive alleles in the homozygous state, while freshwater alleles at other loci underlying physiological traits could potentially have higher dominance coefficients such that they can be expressed in the heterozygous state and homozygous state. Indeed, the lack of phenotypic diversity during the early generations may even provide a basis to allow jackpot carriers and non-jackpot individuals to mate freely, with plasticity in traits like body size reducing the response to selection based on size-assortative mating^71,72^.

Ishikawa et al.^49^ demonstrated that copy number variation at the *fads2* gene involved in docosahexaenoic acid (DHA) synthesis is crucial to colonizing freshwater environments. However, the specimens in the SC2020 sample did not show evidence of elevated *fads2* copy numbers compared to the ancestral Rabbit Slough specimens. This may suggest that the ability to synthesize DHA may be important at later stages of freshwater adaptation, but not the first few generations. Other physiological phenotypes, such as osmoregulation^73,74^, could also be involved in the early stages of freshwater adaptation. For instance, Divino et al. 2016^26^ observed rapid physiological changes in a Scout Lake sample collected in 2012. Our observed lack of differences in morphological phenotypes and the previous studies on physiological traits point to a greater role of the latter in the early stages of rapid adaptation to freshwater environments by oceanic Threespine Stickleback.

## CONCLUSION

Recombination is a fundamental element in the Schluter and Conte transporter model^16^ to describe parallel adaptation by oceanic ecotypes to freshwater environments. It has been clear for a long time that most alleles that encode the differences between freshwater residents and oceanic stickleback exist as standing genetic variants in the oceanic populations^6,10,13,22,46^. In cases of very rapid adaptation, the waiting time for mutation to generate new alleles^75^ should take several thousand years in lake stickleback populations, and even recombination to combine multiple, ancient freshwater-adaptive alleles within individual genomes may take too long. The population would have to rely on nearly intact multiple adaptive alleles that entered the population by a constant gene flow between the oceanic population and other genetically diverging freshwater populations. Using whole genome resequencing of samples from a time series from a rapidly adapting lineage, we showed that recombination played a minimal role at the earliest stages of rapid adaptation in Threespine Stickleback. Instead, rare individuals with large blocks of adaptive standing genetic variants maintained for at least a few generations via reduced recombination will influence whether a new population can adapt quickly to a new environment. Studies of other genomic time series from other Threespine Stickleback will determine how general our results are. Additional studies are required to determine if the adaptive divergence that we observed is characteristic of freshwater colonization by Threespine Stickleback populations and other species. Future research could examine whether F2 jackpot carriers have higher fitness due to their higher survival in freshwater environments or due to the reproductive success of their F1 parents.

## MATERIALS AND METHODS

### Sample collection, library preparation and genome sequencing

All fish used in this study were collected according to an approved protocol from the Institutional Animal Care and Use Committee (IACUC 1446584) at Stony Brook University, as previously noted^12,22^. We extracted DNA samples using the Qiagen DNeasy 96 Blood & Tissue Kit for animal tissue and quantified them with a Qubit Fluorometer 3.0 set to the High Sensitivity option for dsDNA. We used a TN5 transposase-based library preparation using plexWell™ 384 for our low-coverage genomes supplied by SeqWell (Beverly, Massachusetts). We then sequenced the libraries on an Illumina HiseqXten PE 150 bp with an output of approximately 45Gb. We selected 20 high-quality DNA samples for high-coverage resequencing at Beijing Genomics Institute (BGI) using their proprietary DNBseq technology. The target coverage was 30x based on 100bp paired-end sequencing.

### Bioinformatics Processing

Sequencing reads were de-multiplexed into individual genomes based on the i7-barcodes by BGI. The total number of bases sequenced per sample ranged from 0.004 Gb to 1.79 Gb, with a mean of 0.54 Gb. The total number of reads per sample ranged from 21,266 to 11,917,120, with a mean of 3,028,575 reads. After trimming adapters with AdapterRemoval (ver. 2.2.2)^76^, we mapped reads to the Threespine Stickleback genome version gasAcu1-4^12,77^ and stickleback_v5_assembly (https://stickleback.genetics.uga.edu/downloadData/v5_assembly/stickleback_v5_assembly.fa.gz)^78^ using bwa mem -M (v0.7.15-r1140)^79^. Unless stated, we performed all analyses using reads mapped to gasAcu1-4. Libraries sequenced across multiple Illumina lanes and runs were processed separately, and we added read groups using Picard before being merged with samtools ^80^. We also marked duplicates using Picardtools. We performed base recalibration using BaseRecalibrator from GATK version 3.7^81,82^.

### Genotype calling and validation

As mean coverage was ∼1x across samples, we developed a novel method for calling diploid genotype states at freshwater-adaptive loci. As described in the Results, we identified highly linked SNPs within loci that underlie freshwater adaptation, using the correlation coefficient (≥0.99) of SNP allele frequencies from Pool-Seq experiments covering a time series of lake populations undergoing freshwater adaptation: Loberg, Cheney and Scout Lakes^12^. These multi-SNP haplotypes allowed us to call the diploid state of each locus based on an approximate genotype likelihood approach that employed multi-SNP haplotype information. Genotype likelihoods were calculated for individual SNPs within each locus using the approach described in DePristo et al.^83^ and then summed across correlated SNPs based on the freshwater or anadromous allele to obtain approximate likelihoods for anadromous homozygous, heterozygote and freshwater homozygous allelic states for each locus. In addition, we refined this approach by utilizing genotype probabilities for each SNP inferred using an imputation approach (see below). Genotype calls were assigned for a locus if there was a minimum of 3 SNPs with reads with mapping error probabilities less than 0.001 and base call error probabilities less than 0.01. We coded freshwater-adaptive loci that did not meet these filtering criteria as missing.

### Imputation of low-coverage data with Beagle 4.0

We utilized a reference panel of 169 genomes to impute the genotypes of our low coverage TN5 data using beagle 4.0^84^. The reference panel consisted of medium-(approximately 5X) to high-coverage (approximately 30X) published and unpublished genomes from oceanic and freshwater populations. The published genomes were from ^12,85^ (Supplementary Section 7). A list of genomes included in the reference panel and their sources is provided in supplementary table 4.

### Estimating the site frequency spectrum (SFS) with theta and Tajima’s D

To account for our low-coverage genome data, we used RealSFS from ANGSD^86^ to estimate each sample’s site frequency spectrum (SFS) based on genotype likelihoods. We masked ChrXIX and all transposons before estimating the SFS. We used the ancestral genome constructed by Roberts Kingman et al.^12^. Then we imported the estimated SFS into dadi^87^ and folded the SFS, and performed further analyses, including summary statistics like nucleotide diversity (π), Watterson’s estimator of Θ and Tajima’s D^88^. To prevent any bias introduced by unequal sample sizes of various time points, we used dadi to project down each time point to 40 samples.

### Kinship analysis

We utilized two methods to infer pairwise relatedness, both designed for low-coverage data, READv2^33^ and NGSrelate^34^. We performed extensive SNP filtering before analysis for both methods to guard against biases arising from repetitive regions and TN5 library preparation (Supplementary Section 8).

### Inbreeding Estimation

We used NgsRelate^34^ to estimate individual inbreeding coefficients, again using allele frequencies from unrelated specimens from SC2013. To validate the estimates from ngsRelate, we compared runs of homozygosity for our high-coverage SC2020 samples with estimates of inbreeding from ngsRelate for these SC2020 samples (fig. S10). The inbreeding estimates were highly correlated (r=0.86) with the FROH. We also downsampled the high-coverage SC2020 samples and estimated the inbreeding coefficients with ngsRelate. The estimates from the downsampled samples were highly correlated (r=0.822) with the estimates from the high-coverage, although they were underestimated by approximately a factor of 50% (fig. S8).

### Imputation of the non-recombining Y-chromosome (NRY) and haplotype construction

The Y-chromosome will have half the coverage of autosomes, and a considerable fraction of the Threespine Stickleback Y-chromosome is also made of transposable elements ^78^. Therefore, given our low-coverage data, we could not reliably call variants on the Y-chromosome and thereby haplotypes without large amounts of missingness amongst individuals. We therefore developed an imputation approach to allow us to call haplotypes based on the complete linkage of the non-recombining part of the Y-chromosome (NRY), similar to an approach that has previously been used for the imputation of human NRY^35^ (Supplementary Section 10).

### Mitochondrial DNA (mtDNA) haplotype construction

We created a consensus sequence from each sample’s mtDNA using a combination of Samtools view, Bcftools Mpileup, and Bcftools call -c, setting -ploidy 1. We filtered out sites with minor allele frequencies <0.1 and sites with ambiguous nucleotides such as R, Y or K. We excluded non-biallelic sites. After this filtering, we found 10 haplotypes, and we further filtered out haplotypes that occur as singletons at any time, leaving nine haplotypes.

### Estimating copy number variation in fads2 gene

We used GATK’s DepthOfCoverage to calculate the average coverage at the autosomes, sex chromosome (chromosome XIX:12500000-20618466) and the two of the three known copies of the fads2 gene: chromosome XIX:17503349-17507332 and chromosome XII:13777552-13780899. We added the coverage of the fads2 gene copy on chromosome XII and XIX. We also summed the average coverage on the autosomes and the chromosome XIX for all the specimens in the SC2020 sample. We then computed the ratio of the two quantities, with the expectation that a higher copy number of fads2 will lead to an elevated ratio.

### Catch per unit effort estimation

We captured all Threespine Sticklebacks using 1/8 or ¼-inch mesh Gee Minnow Traps set overnight for <24 hours within <5 m of shore and in water <1.5 m deep. We placed the traps in and adjacent to patches of rooted aquatic plants if they had begun to grow. The bottom was mostly sand or silt. We set about 100 traps behind the same three residences on the north shore of the lake. Catch per unit effort (CPUE) is the total number of fish caught divided by “trap hours” The number of trap hours is the product of the number of traps and the number of hours we set.

### Phenotypic characterization

We measured three traits that differ consistently between anadromous and freshwater stickleback^43^ and have diverged rapidly from the ancestral anadromous condition in other recently founded lake populations in the SC2014 sample (Supplementary Section 12). The three traits were lateral plate morphs, gill rakers and standard length.

## ACKNOWLEDGMENTS

This work was supported by NIH R01GM124330 to K.R.V. and M.A.B.,NSF grants DEB-0322818 and DEB-0919184 to M.A.B. and F. J. Rohlf, and the Newcomb College Institute of Tulane University to D.C.H. Many students helped found the Scout Lake population and to capture stickleback from it. We thank them, and particularly P. J. Park, who was instrumental in the fieldwork.

