## supplementary_information_and_figures for "Rare “Jackpot” Individuals Drive Rapid Adaptation in Threespine Stickleback"

1. ESTABLISHMENT, TEMPORAL SAMPLING AND STUDY OF RAPID ADAPTATION IN SCOUT LAKE
2. SEXUAL MATURITY OF SCOUT 2012 AND 2013 SAMPLES
3. DNA EXTRACTION AND QUANTIFICATION
4. CROSSES FROM MILE 87 LAKE FOR VALIDATION OF CALLED GENOTYPES AT FRESHWATER ADAPTIVE LOCI
5. LOW-COVERAGE WHOLE GENOME SEQUENCING OF HUNDREDS OF THREESPINE STICKLEBACK.
6. DETERMINING STATES OF LOCI WITH FRESHWATER-ADAPTIVE ALLELES FROM LOW-COVERAGE SEQUENCE DATA.
7. IMPUTATION OF LOW-COVERAGE SEQUENCES
8. BIOLOGICAL RELATEDNESS
9. INBREEDING
10. IMPUTATION OF THE NON-RECOMBINING Y-CHROMOSOME (NRY)
11. ESTIMATING THE SITE FREQUENCY SPECTRUM (SFS) FOR 4-FOLD AND 0-FOLD DEGENERATE SITES
12. MORPHOLOGICAL PHENOTYPES OF JACKPOT AND NON-JACKPOT CARRIERS IN SC2014
13. ORIGIN OF JACKPOT CARRIERS IN SC2014
14. SUPPLEMENTARY FIGURES

### 1. ESTABLISHMENT, TEMPORAL SAMPLING AND STUDY OF RAPID ADAPTATION IN SCOUT LAKE

In 1990, Threespine Sticklebacks (TS) that resembled anadromous sticklebacks were identified in Loberg Lake (61.559N, 149.262W)[1]. Loberg Lake is a natural kettle lake with a surface area of 4.45 hectares and a mean depth of 5.4 meters in Matanuska-Susitna Borough, Alaska. The lake was previously treated with the broad-spectrum piscicide rotenone to enhance recreational fishing. Threespine Sticklebacks were not detected until 1990. From 1990, Threespine Stickleback samples ranging in size from 59 (1990) to 6500 (1999) were collected from this lake. In 1990, all adults were complete lateral plate morphs, for which anadromous stickleback in Cook Inlet was monomorphic, but some juveniles were low morphs, which are monomorphic in most Cook Inlet lake populations. Thus, the population appeared to be rapidly evolving freshwater phenotypes and annual population sampling was initiated. By 1994, about 45% of the Loberg Lake stickleback sampled were low plate morphs. The frequency of low morphs increased to about 75% in 2001 [1]. Other traits have diverged from the condition of anadromous stickleback toward that of freshwater ones[2,3]. This phenotypic change was one of the earliest cases of rapid phenotypic evolution in Threespine Stickleback[2]. A spring that originates at the south end of the lake and discharges into Spring Creek, in the same drainage as Rabbit Slough, was probably the route by which Loberg Lake was colonized by anadromous stickleback.

Rapid evolution in Loberg Lake and Klepaker's (1993)[4] previous report of rapid evolution in a Norwegian population indicated that oceanic stickleback adapts to freshwater fast enough to study using contemporary time series. To add replicates and greater experimental control, where the source and number of Threespine Stickleback founders are known, we released about 3000 anadromous Threespine Stickleback (TS) from Rabbit Slough (61.536 N, 149.253 W), Matanuska-Susitna Borough, Alaska into each of three lakes, including Scout Lake[5]. Before introducing TS into these lakes, Northern Pike had invaded them, which feeds on small fish, such as TS, causing their extirpation [6]. The Alaska Department of Fish and Game treated these lakes with rotenone to exterminate the invasive Northern Pike. After rotenone treatment, tests for toxicity using caged fish were placed into the lake to determine if they could survive. They treated Scout Lake in the fall of 2009, but it was still too toxic to introduce Threespine Stickleback before the stickleback breeding season ended in June and July of 2010. After further toxicity testing and about 66 trap hours on 11 May 2010, we detected no TS or Northern pike[5]. Thus, we released 3047 anadromous sticklebacks from Rabbit Slough into Scout Lake in June and July 2011[5].

Scout Lake (60.5353N, 150.8322W) is on the Kenai Peninsula, Alaska, USA. It is about 75 m above sea level with a maximum depth of 6.1m. The lake's surface area is about 38.5 ha in a sparsely developed suburban area in Sterling. It is a natural seepage lake without any stream outlets. Aside from Threespine Stickleback, the lake is stocked with coho salmon (*Oncorhynchus kisutch*), Rainbow Trout (*Oncorhynchus mykiss*), and Arctic Grayling (*Thymallus arcticus*).

Samples of Threespine Stickleback have been collected from Scout Lake at least once a year around the start of the breeding season in late May or June. Minnow traps with a mesh of 6.35 mm or 3.175 mm have been set yearly for up to 24 hours at less than 2 m depth and 5 m from shore. After capture, Threespine Sticklebacks are separated from other fish species and transferred to a bucket of lake water to which equal volumes of sodium bicarbonate and MS-222 (tricaine methane-sulfonate) had been added at a high enough concentration to cause the fish to lose equilibrium within 30 seconds and to die within a few minutes. Death was inferred from the failure of the fish to react to tapping the side of the bucket in which they were held and then to pinching the caudal fins of selected fish. We netted the fish out of the bucket, washed them in lake water, and dropped them into 70 % ethanol in deionized water to fill a 1-liter bottle up to about half of the fish and the remainder with ethanol solution. The ethanol was replaced with a fresh 70% ethanol solution within about 24 hours after the lipid from the fish had discolored the ethanol with a yellow hue. The right pectoral fin was usually clipped from the specimen and placed into a small, numbered conical tube of 70% ethanol for DNA extraction, and the remainder of the fish was placed with another fish from the same sample in a numbered, 15-ml tube with 70% ethanol and one fish head up and the other head down.

### 2. SEXUAL MATURITY OF SCOUT 2012 AND 2013 SAMPLES

We removed the gonads of females and males to perform macroscopic inspections of them. Ovaries of females were assigned to six reproductive phases following[7,8]: latent, early maturing, late maturing, mature, ripening, and ripe. These phase assignments have been updated with more recent terminology based upon microscopic histological observations[9] but were used as initially designated. Latent females were considered sexually immature, whereas females in the other five phases were considered sexually mature. Testes of males were assigned to latent or mature phases based on their size and opacity. Latent males had small, transparent or translucent testes, and mature males had enlarged, cloudy to opaque white testes.

### 3. DNA EXTRACTION AND QUANTIFICATION

For each time point, we randomly selected extracts from 96 specimens with high concentrations of DNA(except SC2014, in which there were only 48 specimens) for library preparation with plexWell™ 384 (SeqWell, Beverly, MA, USA). The plexWell™ 384 kit is based on seqWell's proprietary TN5 transposase that involves the insertion of Illumina i7 adapters into individual DNA samples (up to 96 samples), which are then pooled together. An Illumina i5 adapter is inserted into each pool and amplified in a PCR reaction using manufacturer-provided library primer mix and Roche's KAPA HiFi DNA Polymerase readymix. The concentration of each pool was within the manufacturer-recommended range of 4-8ng/μL (RS2019-4.2 ng/μL, SC2013-5.68ng/μL, SC2014-6.84ng/μL, SC2015-4.56ng/μL, SC2017-5.24ng/μL, Crosses-7.56ng/μL). We performed further quality controls by running aliquots of the prepared libraries on 2% agarose gel and an Agilent 2100 Bioanalyzer. We pooled all individuals on a 96-well plate together and added a unique i5-barcoded adapter. Each population of pooled 96 individuals is then sequenced on an Illumina HiseqXten platform, with paired ends read length of 150 bps at an output of around 45Gb. TS has a genome size 460Mb, so 45Gb output yields approximately 100X coverage.

For the high-coverage DNA sequences from SC2020, DNA was extracted using the Qiagen DNeasy 96 Blood & Tissue Kit for animal tissue and was quantified with a Qubit Fluorometer 3.0 set to High Sensitivity option for dsDNA. We selected high quality with minimal evidence of fragmentation to sequence at high coverage. The average concentration of the 20 selected samples was 72.32 ng/uL, ranging from 41.2 ng/uL to 134.8 ng/uL. We sent the DNA for resequencing at the Beijing Genomics Institute (BGI) using their proprietary DNBseq technology.

### 4. CROSSES FROM MILE 87 LAKE FOR VALIDATION OF CALLED GENOTYPES AT FRESHWATER ADAPTIVE LOCI

In early June of 2021, adult sticklebacks were collected from Mile87 Lake (60.914517°, -149.101545°) using unbaited, 10mm mesh minnow traps set within a few meters of the shoreline. Anadromous and freshwater males and females in breeding conditions were transported live in aerated lake water to the temporary laboratory facility in nearby Anchorage, Alaska. We identified ecotype identity based on substantial size and body shape differences between the adults and the anadromous (larger) and freshwater (smaller). We produced four crosses via *in vitro* fertilization: male anadromous x female anadromous, male freshwater x female freshwater, male anadromous x female freshwater, and male freshwater x female anadromous. We euthanized males in an overdose of buffered MS-222, and their testes were dissected and macerated in a drop of embryo medium (2ppt Instant Ocean Sea Salt in distilled water + 1 drop of methylene blue/2L). We stripped ovulated eggs from a gravid female into a Petri dish and subsequently covered them with the extracted sperm. After three minutes to allow fertilization, we rinsed clutches thoroughly, submerged them in fresh embryo medium, and stored them at 7°C. The following day, we separated embryos, removed dead/unfertilized individuals, and refreshed the embryo medium. We maintained clutches each day. Prior to five days post fertilization, clutches were shipped overnight to The College of New Jersey in Ewing, NJ. Embryos began hatching ~12-14 days pf, and fry was reared for an additional three weeks when they were euthanized in an overdose of buffered MS-222 and preserved in 100% ethanol for subsequent DNA extraction. Fish were collected under Alaska Department of Fish and Game permit P-21-006. All procedures were approved by The College of New Jersey Institutional Care and Use of Animals Committee (protocols 1908-001MW1A1 and 2002-001MW1A3).

### 5. LOW-COVERAGE WHOLE GENOME SEQUENCING OF HUNDREDS OF THREESPINE STICKLEBACK.

We previously used pool-seq to produce annual estimates of SNP frequencies in the Scout Lake population from 2012 through 2017 and from a 2009 sample from the ancestral Rabbit Slough population [10] to assess allele frequency changes as the population adapted to freshwater. The genome-wide frequencies of freshwater-adaptive alleles increased rapidly at hundreds of loci, increasing to ~50% within 6 years, along with corresponding changes in freshwater phenotypes[11]. However, while providing extensive information on allele frequencies, pool-seq sacrificed information on individual haplotypes. To examine if haplotypes of freshwater adaptive alleles at Scout Lake re-assembled after introduction to the lake, according to the transporter hypothesis, or were already present in rare large haploblocks, according to the jackpot hypothesis, we used a TN5 transposase-based approach to sequence hundreds of low-coverage, whole genomes from the Scout Lake stickleback time series, including 336 genomes from individuals collected from 2 (n = 96), 3 (n = 17), 4 (n = 47) and 6 years (n = 96) after the Scout Lake population was founded in 2011. Annual samples from these time points are SC2013, SC2014, SC2015, and SC2017. In addition, we sequenced 96 individuals from a 2019 sample from the ancestral Rabbit Slough (RS) population to represent the oceanic fish that were stocked into Scout Lake from the exact location and generated 20 high-coverage (25.7X) genomes from the 2020 Scout Lake sample (SC2020) (Supplementary table 1).

### 6. DETERMINING STATES OF LOCI WITH FRESHWATER-ADAPTIVE ALLELES FROM LOW-COVERAGE SEQUENCE DATA.

We previously identified freshwater-adaptive alleles at 344 loci that experienced rapid and significant frequency increases in three Threespine Stickleback lake populations (including Scout Lake) from Cook Inlet, Alaska[10], which were founded by anadromous stickleback within the last 40 years. Loci with freshwater-adaptive alleles ranged from one or two base pairs to kilobases, with a median size of 27.3 kb[10]. Thus, multiple neighboring SNPs in linkage disequilibrium will tend to share the signal of rapid adaptation for a specific locus. To identify multi-SNP haplotypes for a locus, we selected neighboring SNPs whose freshwater allele frequencies were highly correlated (r> =0.99) with the most significant freshwater-adaptive SNP in the region through our time series. Using this criterion, we obtained a set of 300 loci tagged by multi-SNP haplotypes, each containing three to 3658 SNPs.

We developed an approach that used genotype likelihoods across multiple SNPs to call the diploid state of each locus as homozygous anadromous, heterozygous, or homozygous freshwater from low-coverage genomic data (Methods). We validated the ability of this method to determine the diploid state of the loci by applying it to call haplotypes of a set of experimental crosses between anadromous and freshwater parents from the impoundment at mile 87 along Seward Highway (Supplementary section 3). We observed that estimated states from our likelihood method were highly consistent with expectations from the results of our crosses (fig. S1). Parents were scored as homozygous for the alleles associated at most loci with their ecotype. More importantly, offspring in these experimental crosses inherited states consistent with the states of their two parents. We did not consult that parental genotypes were to call genotypes of their progeny. For example, the offspring of parents with alternative homozygous states at the locus were invariably scored as heterozygous.

We also imputed our low-coverage data with beagle 4.0[12] to enhance genotyping accuracy using a reference panel of 169 medium- to high-coverage whole genomes (Methods). We adjusted our likelihood approach to incorporate imputation-based genotype probabilities rather than individual SNP genotype-likelihoods. The overall pattern of the genotypic state of each locus was not qualitatively different between our likelihood approach and Beagle-imputed genotype probabilities (fig. S1), as the dosage of freshwater-adaptive alleles carried by each individual was similar in both methods. Beagle imputation led to a reduced rate of missingness per locus (280 loci without missingness after vs. 235 before imputation with Beagle).

### 7. IMPUTATION OF LOW-COVERAGE SEQUENCES

Using beagle 4.0 and a reference panel of 169 genomes, we imputed the missing data in the low-coverage sequences. The reference panel comprised genomes mapped to the Gas-Acu1-4 genome, with bwa-mem and base recalibration performed on the bams with GATK's BaseRecalibrator using hard-filtered SNPs from Roberts-Kingman et al.[10]. The coverage of the included genomes ranged from 9 - 62x. It included 69 oceanic fish, 58 from well-established freshwater populations, 40 from experimental transplants, and two of unknown ecotype designation with a global distribution (Supplementary Table 4). We followed GATK's best practices guidelines for variant calling with HaplotypeCaller, joint genotyping, and variant quality score recalibration with VariantRecalibrator. We only included biallelic sites in the reference panel and targeted 6,027,376 high-quality SNPs. We downloaded beagle 4.0 from https://faculty.washington.edu/browning/beagle/beagle.r1399.jar to perform haplotype phasing per chromosome and imputation on all low-coverage data simultaneously. The genotype likelihood used was estimated using GATK's Haplotypecaller. We used recombination maps from ancestral Rabbit Slough, which was previously estimated[10] using LDhelmet[13]. We filtered the recombination maps to exclude absent sites in the reference panel. Then we ran beagle 4.0 as follows java -Xmx90G -jar beagle.27Jan18.7e1.jar gl=chrII.gz ref=WGS_170_recalibrated_Snps6mil_chrII.phased.vcf.gz map=chrII.map impute=True out=All_samples_chrII. After imputation, we combined all chromosomes using bcftools concat.

### 8. BIOLOGICAL RELATEDNESS

We called genotype likelihoods genome-wide for all SNPs identified previously by Roberts Kingman et al.[10] using GATK Unified Genotyper v3.7.0[14]. We estimated allele frequencies for each Scout Lake sample separately using the approach of Kim et al.[15]. We then intersected these allele frequencies with allele frequencies estimated from pool-seq data from Kingman et al.[10] from the same time points using the approach of Lynch et al.[16]. For each time point, any SNP that had a total depth of coverage (across all samples) of less than half or double the mean coverage for both the TN5 and pool-seq data was included in the genome mask. In addition, we performed Fisher's exact test on a 2x2 contingency table containing the depth of each allele in the TN5 and Pool-Seq data and added any SNPs with a p-value less than 0.05 to the mask. We merged masks from each time point, known repeats, and transposable elements. Then, we applied this mask to all time points, resulting in a set of 2,119,671 high-quality SNPs for relatedness analysis. We made haplodiploid genotypes for use in READv2 based on the highest allele depth[17,18], which estimates the degree of relatedness between pairs of individuals by using the proportion of allele mismatches across the genome. We used READv2[18], which estimates up to third-degree relatedness. We validated READv2 results using ngsRelate[19], which utilizes population allele frequencies to estimate identity-by-descent probability. We ran ngsRelate with options ngsRelate -F 1 -h input.vcf -O output -z samp_list -A AF, where AF is the allele frequency estimated from a set of unrelated samples from SC2013 (again estimated using the approach of Kim et al.[15] )

We used Cotterman coefficients[20] for first-degree relatives to distinguish parent-offspring from sibling pairs. We relied on the Cotterman coefficients estimated from ngsRelate instead of READv2. When there is no inbreeding, the Jacquard coefficients J9, J8 and J7 map to the Cotterman coefficients. Based on the Cotterman coefficients, the five first-degree relatives observed in our Scout Lake samples were likely siblings. However, READ2 designates 4 out of these five as parent and offspring. In READv2, the Cotterman coefficients k0 and k2 correspond to the proportions of windows classified as unrelated and identical. This proportion is expected to be low for parent-offspring pairs and higher for siblings.

### 9. INBREEDING

To validate the estimates of inbreeding coefficients from ngsRelate (fig. 2D), which is designed for low-coverage datasets, we used PLINK to estimate the runs of homozygosity for our high-coverage SC2020 samples. We compared the results with estimates of inbreeding from ngsRelate for them. We estimated runs of homozygosity by first performing LD pruning in PLINK with options `--indep-pairwise 50 5 0.5`. We then estimated the runs of homozygosity with PLINK, option `--homozyg --homozyg-window-kb 5000`. There is a high correlation (r=0.86) between the runs of homozygosity and estimated inbreeding coefficients from ngsRelate (fig. S8).

We also downsampled the high-coverage SC2020 samples to 1X coverage using `samtools` (v1.9) with option `samtools view -s x -b input.bam -o downsampled.bam`, where x is calculated as 1/y, and y is the coverage of the high-coverage sample estimated using GATK's `DepthOfCoverage`. We used ngsRelate to estimate the inbreeding coefficient using allele frequencies from SC2013. There is a high correlation (r=0.822) between estimates from downsampled and high-coverage samples (fig. S13)

### 10. IMPUTATION OF THE NON-RECOMBINING Y-CHROMOSOME (NRY)

To impute the NRY, we identified males based on the coverage of ChrXIX and the autosomes. In males, we expect the coverage on the ChrXIX to be half that of autosomes. We identified 258 males. We mapped our reads to the v5 of the Threespine Stickleback genome (https://stickleback.genetics.uga.edu/downloadData/v5.0.1_assembly/stickleback_v5.0.1_assembly.fa.gz ) [21] and verified males with the coverage on the chrY. We then estimated the average coverage for sites on the Y-chromosome of all the identified males. The mean coverage on the Y-chromosome across all males was 0.728 (range: 0.09, 1.94). We divided the Y-chromosome into overlapping tiles of 10 Kbp, used a step size of 500bp, and calculated the average coverage for each window. Then, we filtered out windows that did not fall within half the average ± 1 SD of the autosome coverage (0.94). We combined contiguous windows into one interval and found sites within these intervals. We ended up with 8,541,267 sites after this filtering. We used GATK's `Haplotypecaller` to call SNPs from these filtered sites, setting ploidy to 1. There were 58,685 SNPs.

We filtered further by removing indels and SNPs with >2 alleles, leaving 29,300 biallelic SNPs. For the remaining SNPs, if the genotype was called, we checked if the allelic read depth was equal to the genome depth. We then include only SNPs with minor allele frequencies greater than 0.1. We only included sites with a genotype in at least 100 of the 258 male individuals. After this filtering, there were 2102 SNPs, which we used to compute the correlation matrix used in the imputation described in the Methods section of the main text and described fully below. There was an average missingness rate of 56% per site, and each sample had between 788 and 2094 sites with missing data out of 2102 sites resulting after these filtering (37% to 99% missing rate).

Our approach for imputation relies on the expectation that haplotypes have had the same genetic background due to the complete linkage of NRY for a long time (assuming an infinite site model). Therefore, we can use sites with highly correlated allelic frequencies to predict one another. A similar approach was used previously to impute human NRY[22]; however, note that Lippold et al. [22] defined haplotypes before imputation using sites without missing genotypes as a reference haplotype set, which were then used for imputing missing sites. The coverage of our data on the stickleback NRY does not allow us to make such a reference set. Although we could use high coverage data to construct such a reference set, this will require hundreds to thousands of genomes, especially in our case, where we sought haplotypes that might be rare and are unlikely to be included in reference sets from a few hundred genomes.

We computed an $n \times n$ correlation matrix, where $n$ is the number of sites. For each site, we find other sites whose allele frequencies are highly correlated with it and use them to predict the missing genotypes at that site. This approach scales up the effective coverage for each site, such that if a site with the original estimated coverage of 0.2X has 10 other sites predicting it, in principle, the coverage will now be 2X. Because sites can be correlated due to being in the same read, we only use predictive sites that are 10Kbp from the predicted site.

After imputation, we implemented a greedy algorithm to select sites with the least missingness to construct Y-haplotypes. We set the correlation cut-off at 0.95 and a minimum call rate (i.e., the fraction of individuals with genotypes after imputation) to 0.8. We performed various imputation schemes by varying these parameters, such as correlation coefficient and rate of non-missingness post-imputation, but this did not affect the pattern in our data.

### 11. ESTIMATING THE SITE FREQUENCY SPECTRUM (SFS) FOR 4-FOLD AND 0-FOLD DEGENERATE SITES

We estimated the SFS for all time points using ANGSD[23]. For SC2015, we identified four duplicates through our relatedness analyses (supplementary section 8), which were removed before estimating the SFS. We used degenotate (https://github.com/harvardinformatics/degenotate) to compute the degeneracy of coding sites across the stickleback genome. In total, we identified 24,891,530 0-fold and 6,348,969 4-fold degenerate sites. Then, we used the -sites option from ANGSD to estimate the SFS for the 0-fold and 4-fold sites. We used realSFS from ANGSD to estimate the SFS for each timepoint and used dadi[24] to fold the SFS.

### 12. MORPHOLOGICAL PHENOTYPES OF JACKPOT AND NON-JACKPOT CARRIERS IN SC2014

Specimens for morphological analysis were fixed in 10% buffered formalin, transferred to 50% isopropyl alcohol, and stained in Alizarin Red S, as described in Bell et al. (2004) or Aguirre et al. (2008)[1,25]. We scored three traits, lateral plate morphs and numbers of gill rakers and dorsal fin rays, which differ consistently between anadromous and freshwater stickleback[26] and have diverged rapidly from the ancestral anadromous condition in other lake populations that were found recently by anadromous stickleback[1,4,27].

Lateral plates are enlarged scales that form a single row along each body side and can be scored with the naked eye. There are three major lateral plate morphs (e.g.,[28,29]) that are strongly influenced by the *Eda* gene[28,30,31]. The complete morph is ancestral to the other two morphs [30]. Anadromous Threespine Stickleback is usually complete morphs[32,33], and the Rabbit Slough population is monomorphic complete[25]. Complete morphs have a plate on each body segment; specimens typically have about 33 plates per side. Complete morphs may be homozygous for the ancestral anadromous allele or heterozygous for it and the derived freshwater allele[30,31,34]. Low morphs usually have four to seven plates per side but no more than 10, which are restricted to the anterior third of the body. Lows are homozygotes for the freshwater allele of *Eda*. Partial morphs have more than 10 anterior plates, an unplated area, and a separate row on the caudal peduncle[28]. They are heterozygous for *Eda*.

Fin rays are jointed bones that support the fins of bony fishes. All fin rays in the dorsal fin were counted under a dissecting microscope. Gill rakers were counted by slitting the membrane at the ventral end of the operculum, and the operculum was lifted to count the number of gill rakers on the anterior edge of the first right gill arch whether they were ossified or not.

### 13. ORIGIN OF JACKPOT CARRIERS IN SC2014

We sought to rule out the unlikely possibility that some resident Threespine Stickleback in Scout Lake survived the rotenone treatment in 2009 and mated with the introduced anadromous stickleback in 2011 to produce the jackpot carriers captured in 2014. Since the previous Threespine Stickleback that inhabited the Scout Lake likely had genotypes similar to nearby established freshwater populations, we retrieved all 51 genomes that were marked freshwater from the Pacific Northeast from ref [10] as a proxy for the previous residents of Scout Lake (see supplementary table 7 for complete list of genome names). We also used 20 genomes of individuals collected from Rabbit Slough in 2009 to represent the anadromous population that was used to found the lake (fig. S15a). We then applied our approximate likelihood-based approach (described in the main text and supplementary section 6) to call genotypes of freshwater adaptive loci for all the 51 freshwater-marked genomes (fig. S15b). Out of the 51 genomes, we found two with predominantly marine genotypes at these loci despite being marked freshwater. Therefore, we applied a threshold and filtered out all individuals with less than 30% freshwater content at these loci (the minimum proportion of freshwater content we observed in individuals sampled in SC2020). After this filtering, 46 genomes remained. We then created hybrid genotypes from the remaining individuals from the Pacific freshwater populations and the Rabbit Slough individuals and estimated the genotype of the loci (fig. S16). We found that the individuals had 50 to 63% freshwater content at loci (fig. S17) compared to the 12.5% to 49.5% content in the individuals we sampled in SC2014. This result suggests that the jackpot carriers sampled in SC2014 were not hybrids from remnant resident freshwater stickleback that might have survived the rotenone treatment. Instead, given that previous studies have identified jackpot carriers in anadromous populations [10,35], jackpot carriers represented in the Scout Lake were descendants of anadromous jackpot carriers from Rabbit Slough.

14. SUPPLEMENTARY FIGURES

Link to pdf: Supplementary figure 1

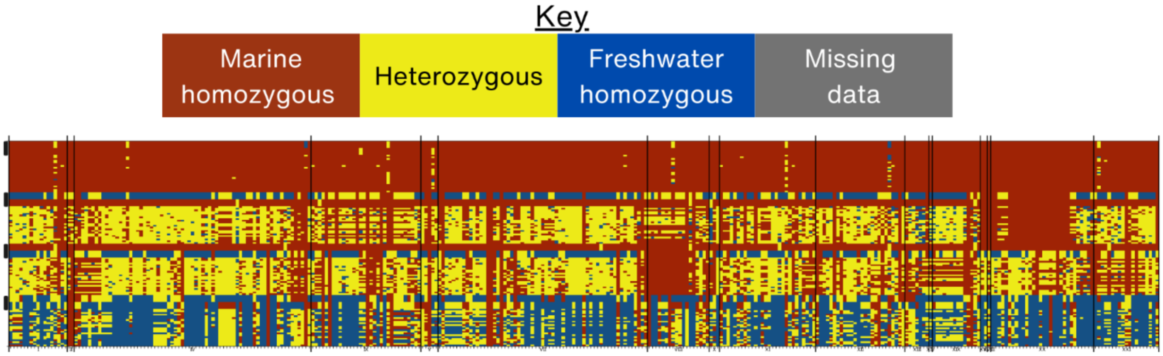


S1. **Haplotypes of crosses used for validating our genotype calling**. The crosses were made using stickleback collected from a lake at mile 87 of Seward Highway a few kilometers east of Girdwood, Alaska. The black rectangles indicate the genomes from parents used for the crosses. The topmost parents are two marine parents followed by their 22 offspring. The next two parents include one marine and one freshwater (one with the male being freshwater and female being marine and vice

Link to pdf: Supplementary figure 2

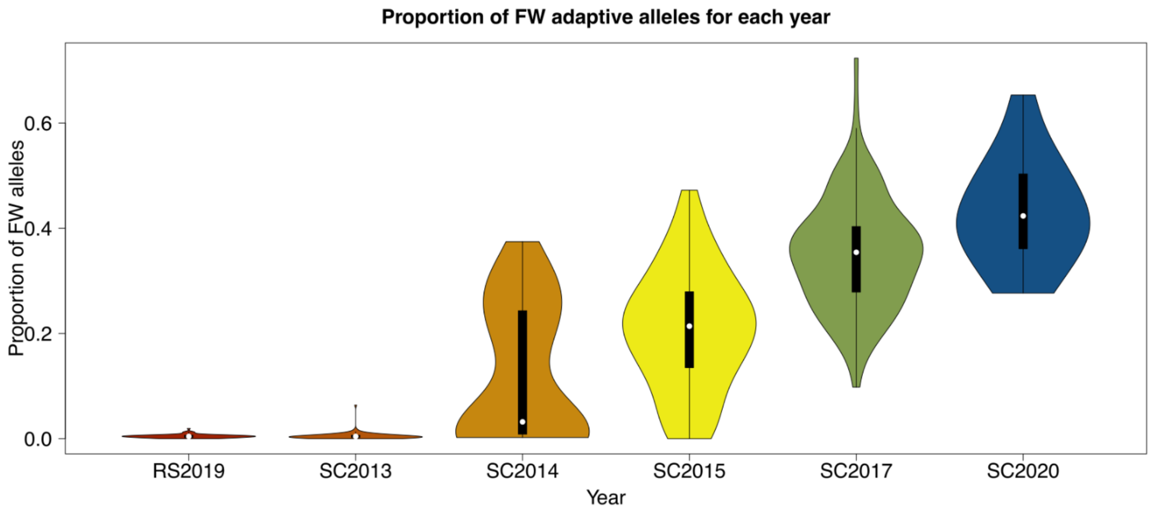


S2: **Quantification of the freshwater content for each sampled timepoint.** The proportion of freshwater adaptive alleles for each time point was estimated by counting the number of freshwater alleles at called loci (out of 344 adaptive loci identified by Roberts Kingman et al. (see supplementary section 6)

Link to pdf: Supplementary figure 3

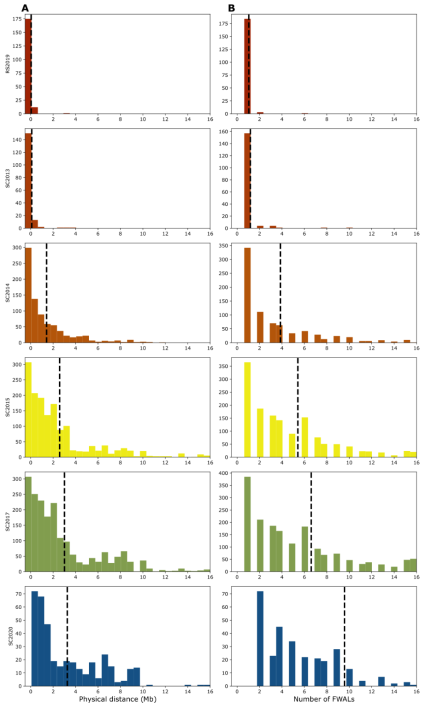


S3 **Physical distance between freshwater adaptive loci and the number of loci that form a contiguous block at various time points during early**

Link to pdf: Supplementary figure 4

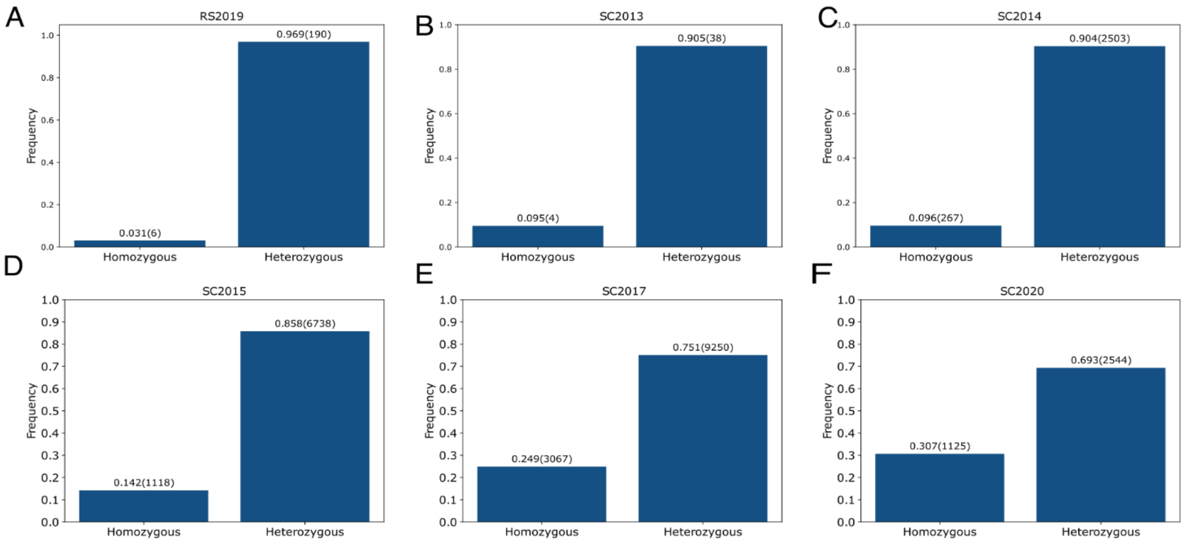


S4 **Heterozygosity at contiguous blocks of loci involved in freshwater adaptation across the various timepoints**

Link to pdf: Supplementary figure 5

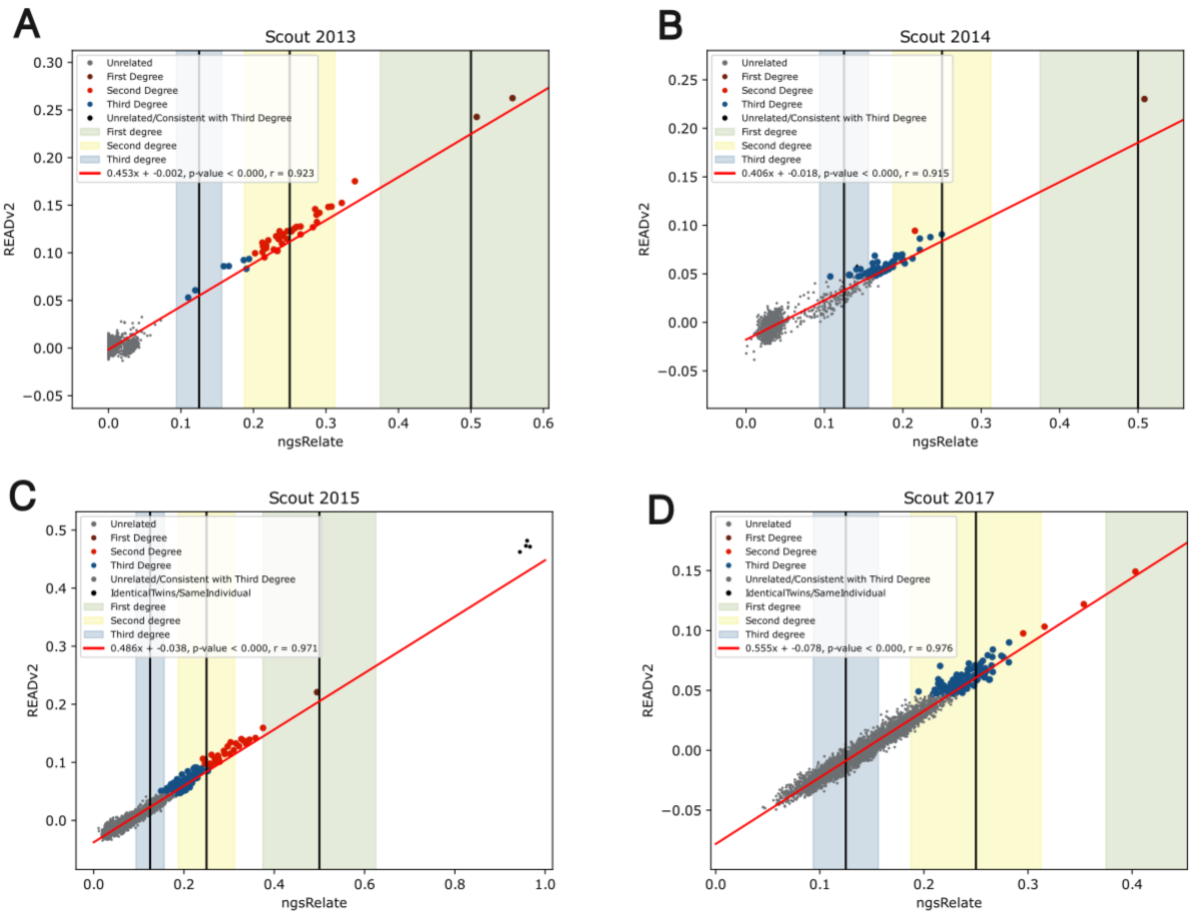


**S5: Relatedness estimates between READv2 and ngRrelate.** The relatedness between all the Scout Lake samples (SC2013, SC2014, SC2015, SC2017 and SC2020) estimated with READv2(Alaçamlı et al. 2024) and ngsRelate(Hanghøj et al. 2019) using SNPs filtered as described in section S8.

Link to pdf: Supplementary figure 6

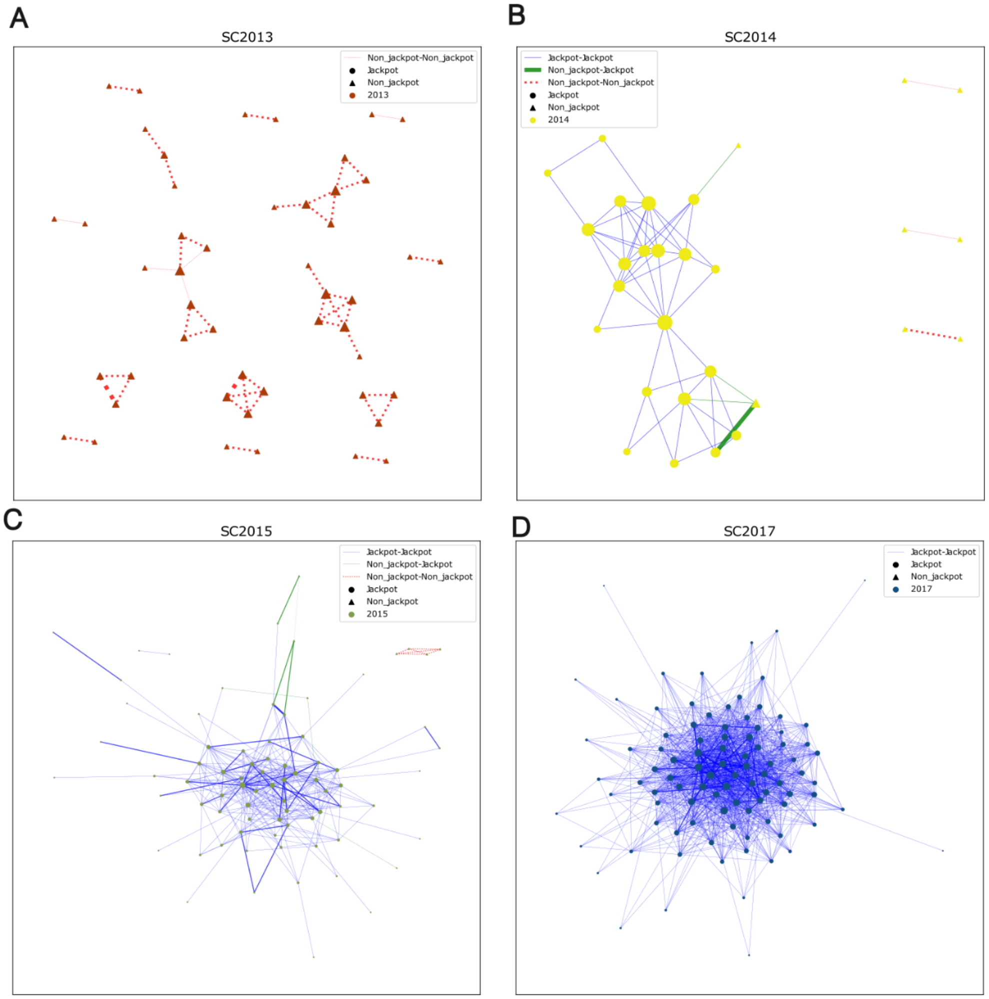


S6 I**ntra-year biological relatedness:** **(A)** Relatedness between specimens sampled in SC2013. **(B)** Relatedness between specimens sampled in SC2014. **(C)** Relatedness between specimens sampled in SC2015. **(D)** Relatedness between specimens sampled in SC2017. The thickness of edge reflect the degree of relatedness between the: the thickest edge first-degree, intermediate second-degree and the thinnest edge third-degree relatives. The size of node increases with increasing number of relatives.

Link to pdf: Supplementary figure 7

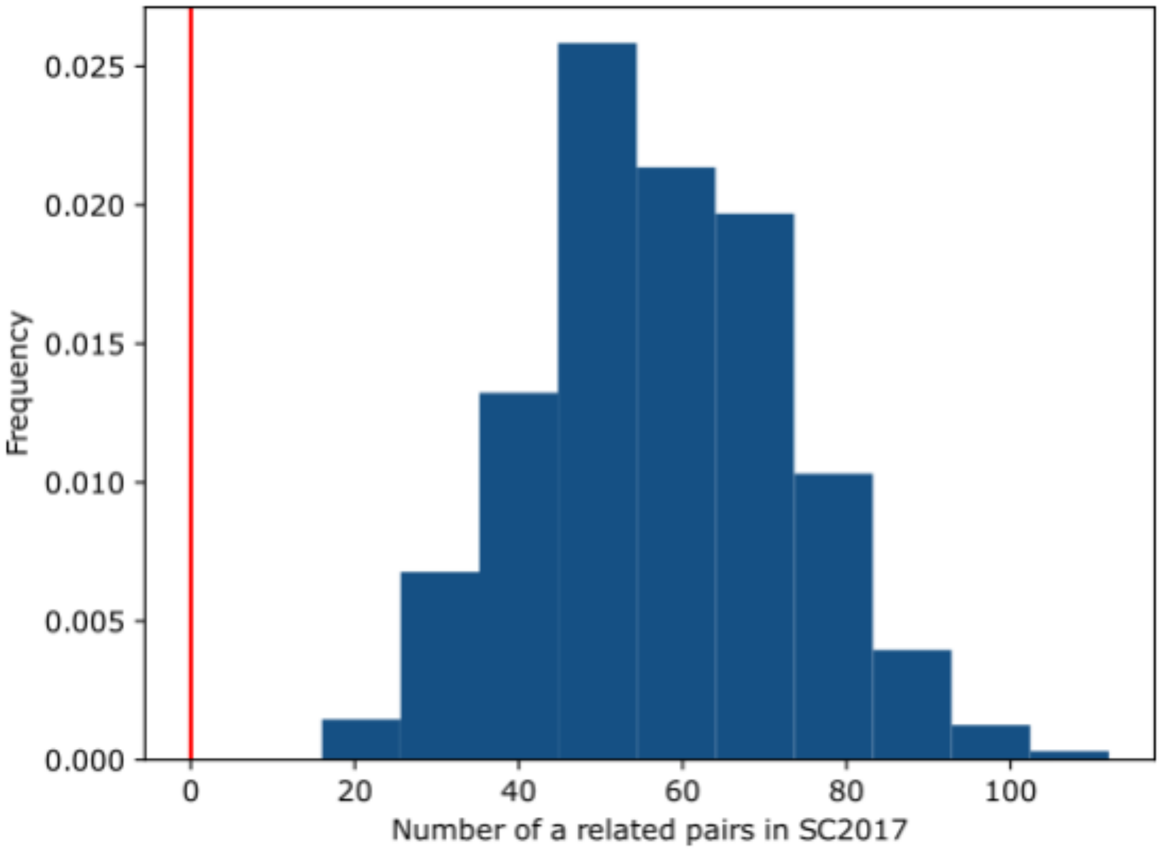


**S7: Permutation of relatedness among SC2017:** We randomly sampled 20 individuals from the 96 SC2017 sample 1000 times and observed the number of pairwise relatives of at least third degree. The red vertical line indicates the observed number of relatedness in the SC2020 sample.

Link to pdf: Supplementary figure 8

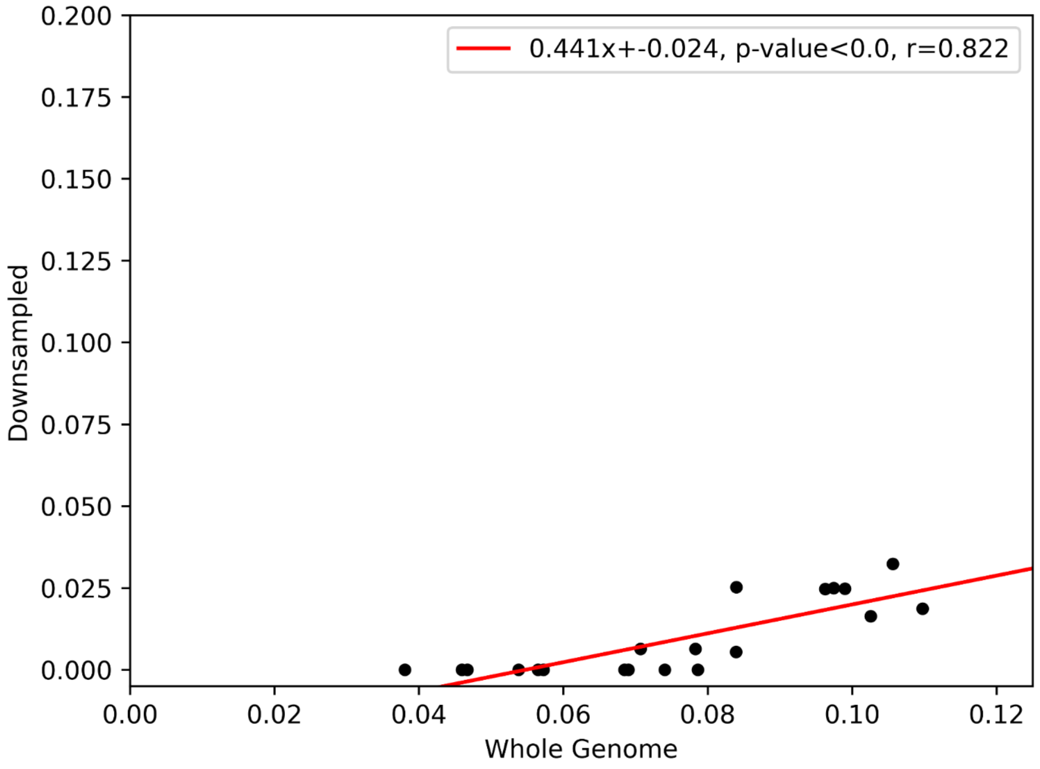


S8 I**nbreeding coefficients estimated from high coverage whole genomes and their downsampled low coverage:** We used ngsRelate to estimate the inbreeding coefficient. There is a high correlation (r=0.822) between estimates from downsampled and high coverage specimen

Link to pdf: Supplementary figure 9

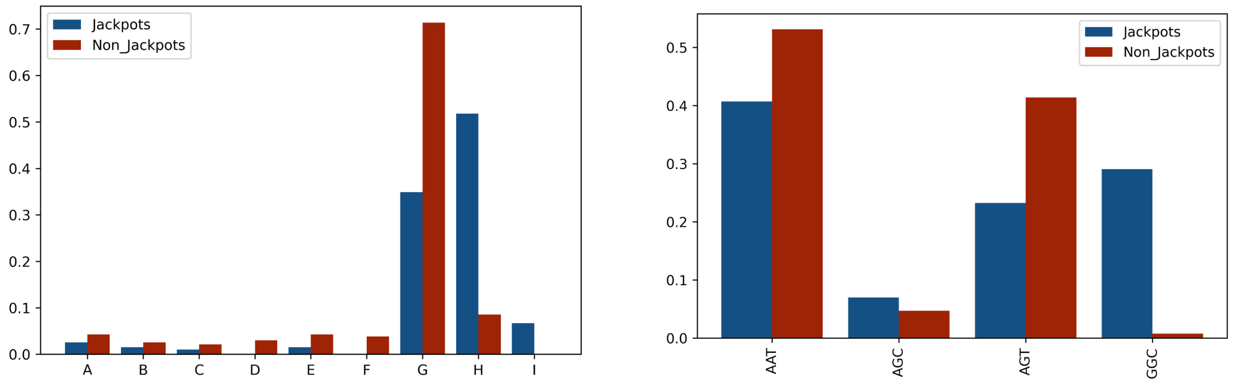


S9 **Frequencies of various haplotypes in jackpots and non-jackpots** (A) Frequencies at the mitochondrial DNA and (B) Frequencies at the Y-chromosome. There are nine

Link to pdf: Supplementary figure 10

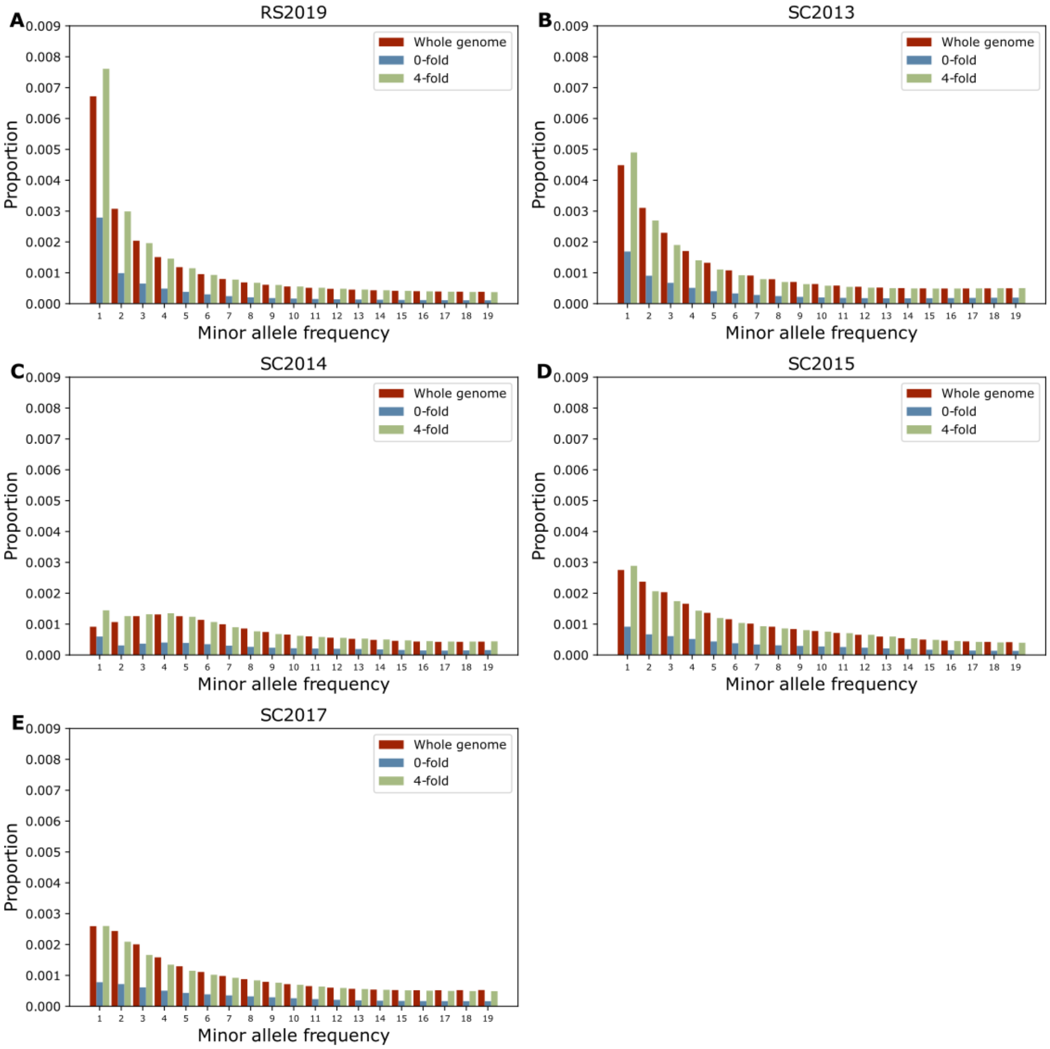


S10: **Site Frequency spectrum at the whole genome, 0- and 4- fold sites for all sampled timepoints except SC2020.**

Link to pdf: Supplementary figure 11

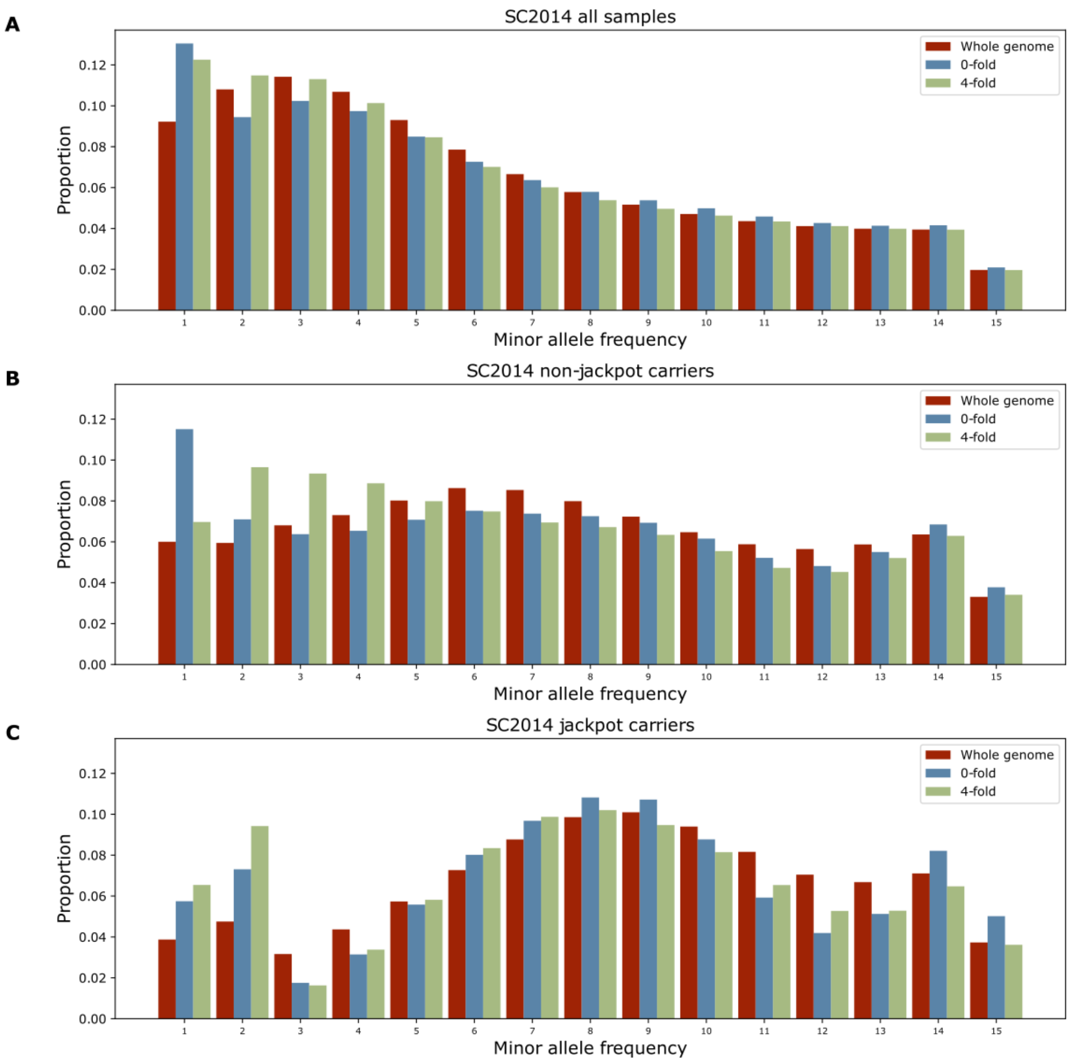


S11 **Site frequency spectrum of jackpot and non jackpot carriers found in SC2014 at the whole genome, 0-fold and 4-fold sites.**

Link to pdf: Supplementary figure 12

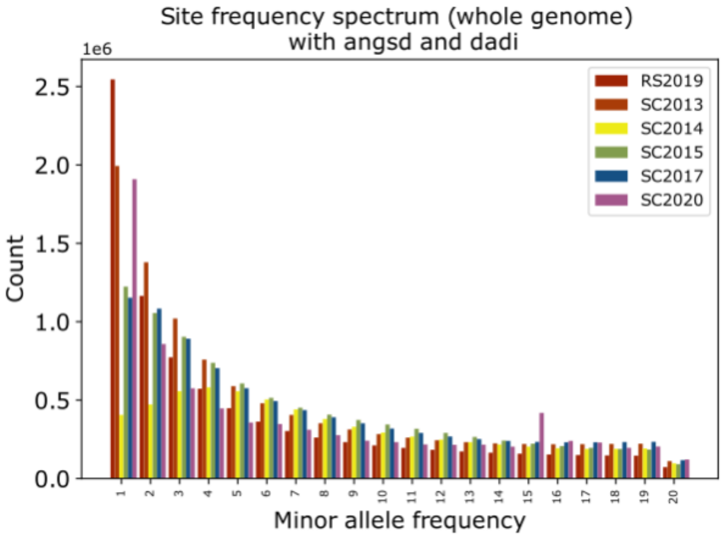


S12. **Site Frequency spectrum (SFS) estimated from the whole genome for all timepoints sampled**. The SFS was folded in dadi and projected down to 40. For clarity, we only show the first 20 polymorphic sites in the SFS.

Link to pdf: Supplementary figure 13

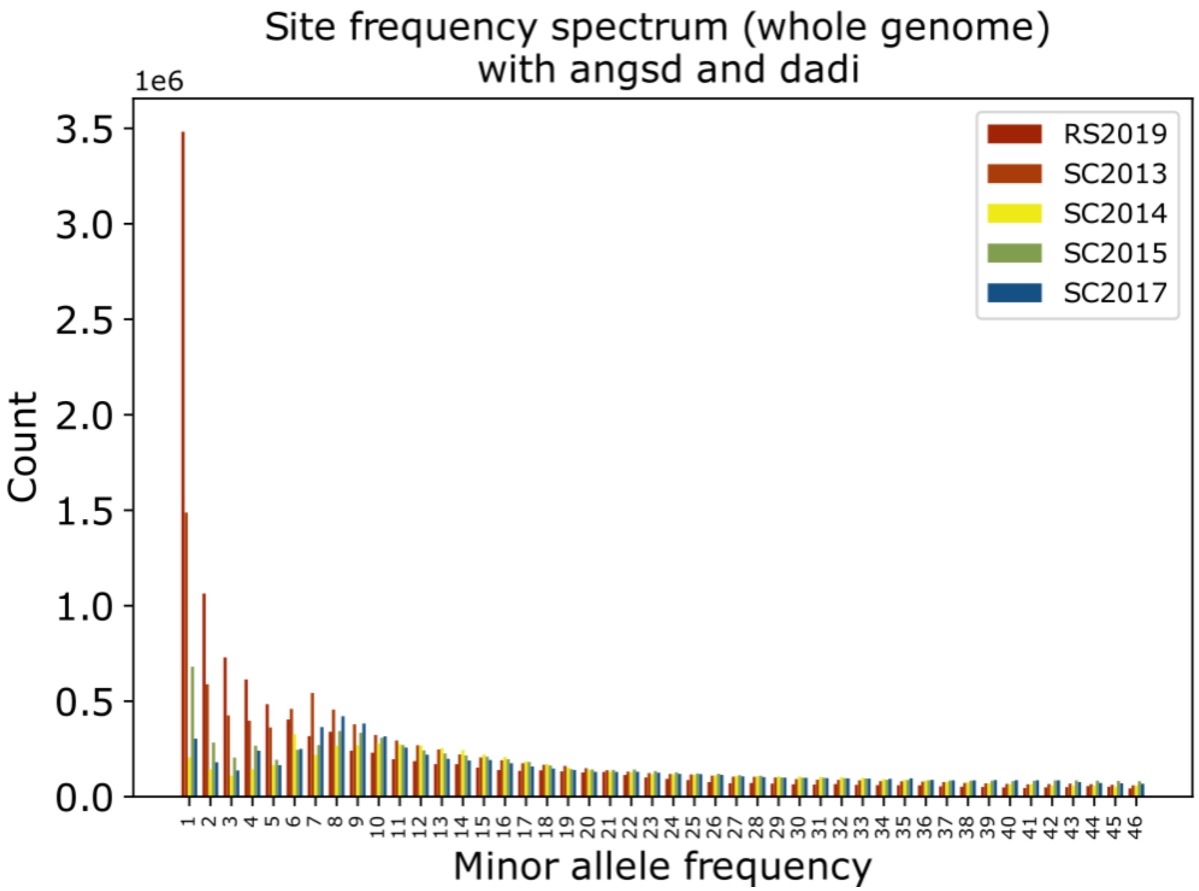


**S13: Raw Site Frequency spectrum (SFS) estimated from the whole genome for all timepoints sampled except 2020**. The SFS was folded in dadi but not projected down to 40.

Link to pdf: Supplementary figure 14

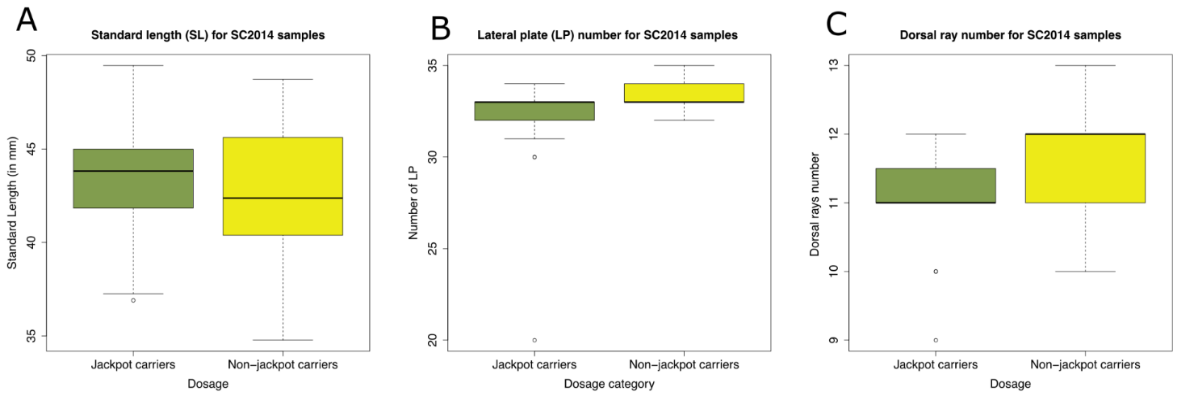


S14 **Morphological phenotypes measured for jackpot carriers and non-jackpot carriers in SC2014**

Link to pdf: Supplementary figure 15

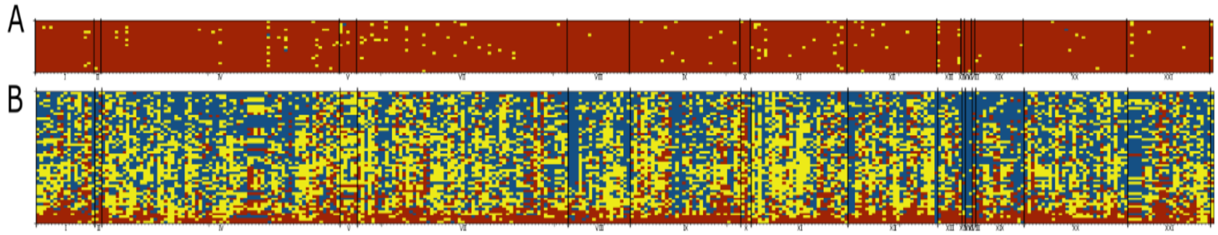


S15 **Genotypes of specimen described in section S13** (A) Rabbit Slough genomes collected in 2009 (B) Genomes from freshwater populations in the Pacific from Roberts Kingman et al.

Link to pdf: Supplementary figure 16

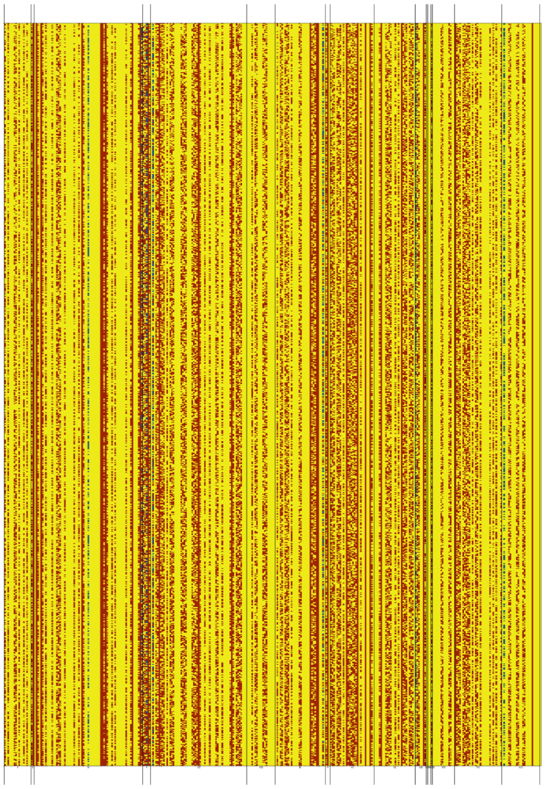

S16 **Hybrids generated by oceanic individuals and individuals from freshwater**

Link to pdf: Supplementary figure 17

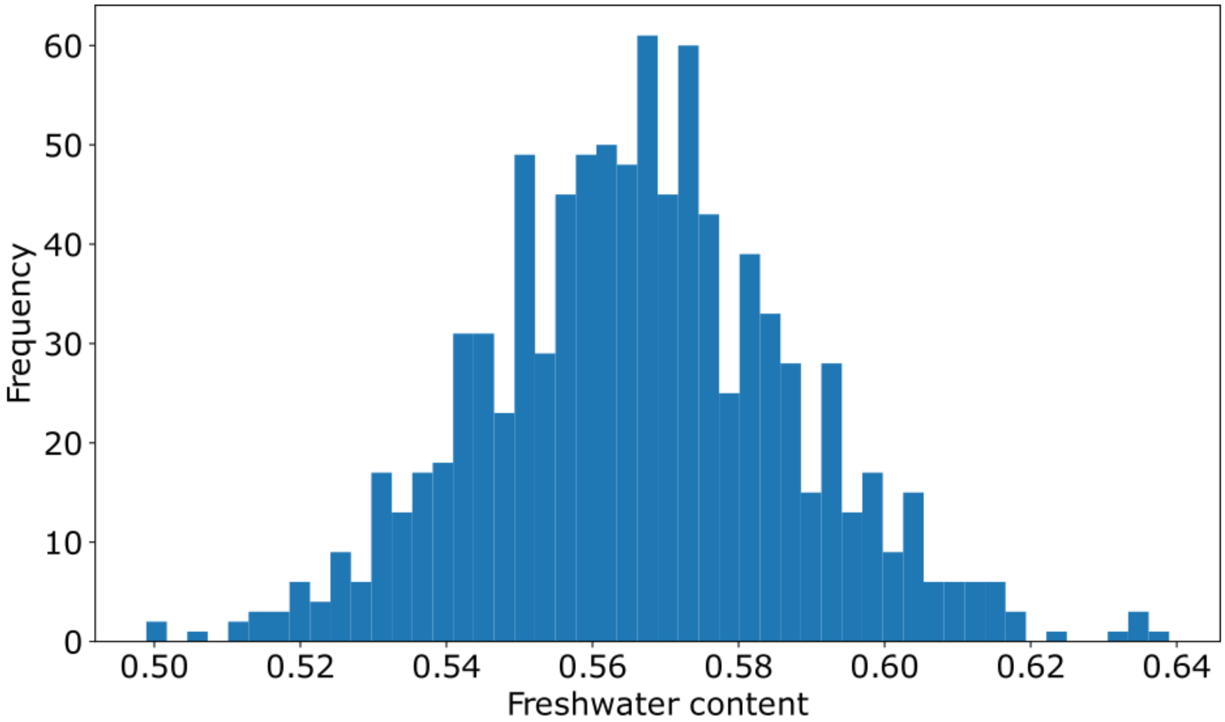


S17 **Distribution of freshwater content of hybrids of oceanic individuals and freshwater individuals**
